# A collaborative resource platform for non-human primate neuroimaging

**DOI:** 10.1101/2020.07.31.230185

**Authors:** Adam Messinger, Nikoloz Sirmpilatze, Katja Heuer, Kep Kee Loh, Rogier B. Mars, Julien Sein, Ting Xu, Daniel Glen, Benjamin Jung, Jakob Seidlitz, Paul Taylor, Roberto Toro, Eduardo A. Garza-Villarreal, Caleb Sponheim, Xindi Wang, R. Austin Benn, Bastien Cagna, Rakshit Dadarwal, Henry C. Evrard, Pamela Garcia-Saldivar, Steven Giavasis, Renée Hartig, Claude Lepage, Cirong Liu, Piotr Majka, Hugo Merchant, Michael P. Milham, Marcello G.P. Rosa, Jordy Tasserie, Lynn Uhrig, Daniel S. Margulies, P. Christiaan Klink

## Abstract

Neuroimaging non-human primates (NHPs) is a growing, yet highly specialized field of neuroscience. Resources that were primarily developed for human neuroimaging often need to be significantly adapted for use with NHPs or other animals, which has led to an abundance of custom, in-house solutions. In recent years, the global NHP neuroimaging community has made significant efforts to transform the field towards more open and collaborative practices. Here we present the PRIMatE Resource Exchange (PRIME-RE), a new collaborative online platform for NHP neuroimaging. PRIME-RE is a dynamic community-driven hub for the exchange of practical knowledge, specialized analytical tools, and open data repositories, specifically related to NHP neuroimaging. PRIME-RE caters to both researchers and developers who are either new to the field, looking to stay abreast of the latest developments, or seeking to collaboratively advance the field.

## 1. Introduction

When navigating an unfamiliar city, people orient themselves based on a few landmarks, use a map for detailed navigation, and take advice from locals and previous visitors about where to go and which areas to avoid. Navigating a highly specialized research field, such as non-human primate (NHP) neuroimaging, is conceptually very similar. In order to orient, access to basic information and support from knowledgeable peers is invaluable. Here we present the **PRIMatE Resource Exchange** (PRIME-RE, https://prime-re.github.io), a new community-driven platform that promotes communication among peers and facilitates the exchange of specialized knowledge and resources related to NHP neuroimaging.

Translational neuroimaging in animals bridges the gap between non-invasive imaging studies in humans and invasive neural recordings that are only carried out in animal models (and occasionally in neurosurgical patients). The approach is pivotal for our understanding of the neuronal basis of neuroimaging signals (Logothetis, 2003; Logothetis et al., 2001; Logothetis and Wandell, 2004), guides the selection of recording targets for invasive studies, and reveals the broader brain networks involved in particular cognitive functions or brain processes. Cross-species comparison of neuroimaging results can furthermore provide insight into the evolution of the brain and its capabilities (Friedrich et al., this issue; Heuer et al., 2019). Finally, combining neuroimaging in animals with invasive methods that influence neuronal activity makes it possible to draw causal inferences about brain function and test potential human therapies (Klink et al., this issue).

The NHP brain is similar to that of humans in many respects. Due to the evolutionary proximity of NHPs to humans, their ability to perform complex cognitive tasks, and the vast body of existing neuronal knowledge from invasive NHP studies, NHP neuroimaging is a crucial branch of translational imaging (Chen et al., 2012; Farivar and Vanduffel, 2014; Logothetis et al., 1999). However, the standard hardware and software used in human neuroimaging is generally not directly suitable for NHP neuroimaging, which requires highly specialized approaches. As a result, most research groups have developed their own in-house solutions that often require substantial further development for compatibility with other sites. These custom solutions often remain in-house and lack extensive documentation, which is far from ideal for peer review, collaboration, best practices, and reproducible science. In recent years, the international NHP neuroimaging community has begun embracing a culture of data-sharing, open science, and collaboration. These efforts led to the PRIMatE neuroimaging Data Exchange initiative (Milham et al., 2018), a global workshop to discuss strategies for the future of the field (Milham et al., 2020), and this special issue of NeuroImage on NHP neuroimaging.

Once NHP neuroimaging data are collected or shared, a new set of challenges presents itself. Most neuroimaging software packages have been developed for use in humans and cannot be easily applied to NHP data due to built-in assumptions, such as a larger field-of-view, standardized voxel size, or a higher contrast between gray and white matter. The large number of choices involved in neuroimaging data analysis is a hotly-debated issue in human neuroimaging (Botvinik-Nezer et al., 2020). For NHP neuroimaging, these issues may be even more pressing due to the custom-solutions that are often required to make existing software compatible with NHP data. The field of human neuroimaging has started to address these issues with initiatives for standardized, reproducible analysis pipelines (BIDS, Nipype, etc) (Gorgolewski et al., 2011, 2016), data sharing (OpenfMRI, Zenodo, Neurovault, etc) (Gorgolewski et al., 2015; Poldrack et al., 2013), and collaborative and open coding (GitHub, GitLab, etc). NHP neuroimaging has been lagging behind in adopting such collaborative open science initiatives, but people’s willingness to share and be open about the resource solutions they developed is steadily increasing (Balbastre et al., 2017; Tasserie et al., 2020) and new developments in information technology promote these initiatives, while creating avenues to assign explicit credit to developers. The global NHP neuroimaging consortium (Milham et al., 2020, 2018) aims to facilitate this progress with the introduction and curation of the **PRIMatE Resource Exchange** (PRIME-RE, https://prime-re.github.io), a community-driven online platform for the exchange of knowledge and resources concerning the acquisition, analysis, and visualization of NHP neuroimaging data.

PRIME-RE acts as an open hub where researchers from all countries and career stages can find and contribute information and resources related to the challenges and advantages of NHP neuroimaging. Dynamic collections of software tools are grouped by analysis-category to provide an overview of existing solutions to common challenges. Links to communication outlets allow researchers to meet international colleagues, discuss challenges and solutions, and start new collaborations. Resources are maintained by their respective developers, with PRIME-RE serving as an evolving access point with descriptions of the tools, their software requirements, a list of authors, relevant citations, and links to the tools themselves.

## 2. Methods

This Methods section will offer a brief description of the main structure of the PRIME-RE platform, including its main components, methods for people to contribute and interact, and possibilities for future growth. The lightweight PRIME-RE website (https://prime-re.github.io/) is intended to be dynamic and continuously evolve based on input from the research community. It can serve both as a starting point for researchers that are new to the field of NHP neuroimaging and as a meeting ground for more experienced researchers in the field. This initiative emerged from the Global Collaborative Workshop 2019 of the PRIME-DE consortium and subsequent BrainHack (Milham et al., 2020) and has since attracted a critical mass of active contributors dedicated to open and collaborative NHP neuroimaging. The platform is hosted on GitHub, which greatly facilitates community-driven contributions while also maintaining a complete version history of its content and evolution.

Content-wise, PRIME-RE does not aim to host, curate and maintain all available tools and documentation regarding NHP neuroimaging. Instead, it chooses an agile approach with standardized listings of resources, categorized by the type of challenge they address. These listings include a brief description, developer contact information, and a link to the developer-maintained resource. This model furthermore ensures that the most recent version of a resource will be found at its source and that developers receive credit for their efforts. The standardized listings also make it easy for developers to submit their resource for inclusion on PRIME-RE through a pre-formatted submission form in which they answer a few basic questions about their resource (Table 1). These answers form the basis of a resource’s listing on PRIME-RE. If the developers are already hosting their resource online, a submission to PRIME-RE can be completed in a few minutes.

**Table 1.**
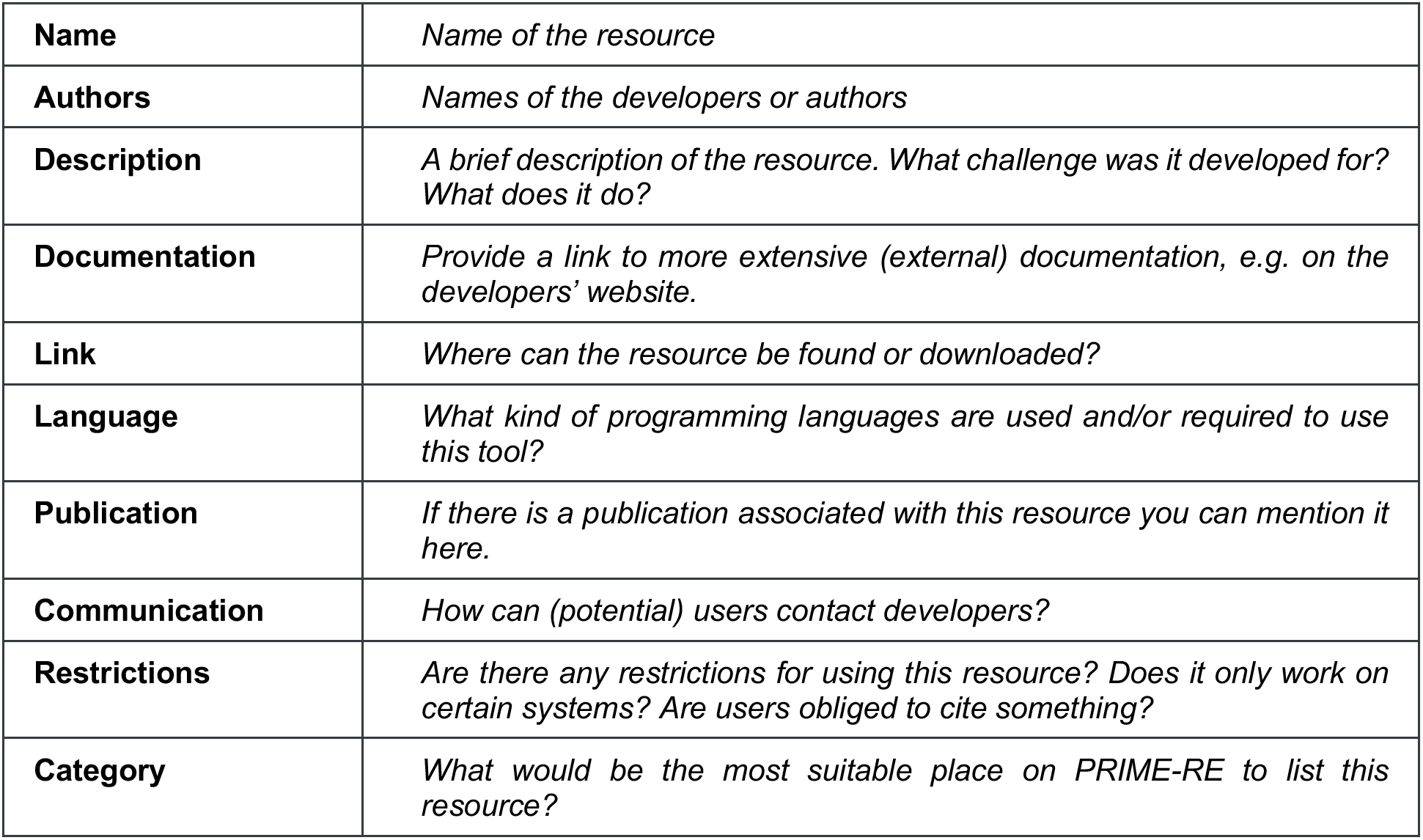
Standardized information table for the submission of a resource for inclusion on PRIME-RE. When developers click a ‘Contribute’ button they are taken to a pre-formatted form where they can provide this information. Any member of the community with access to the website’s backend can then include the new resource under the correct website category. In addition to this table, the submission form also has checkboxes with which the submitter can confirm that resource development occurred in accordance with all applicable directives and guidelines.

Submissions to PRIME-RE are approved for inclusion through community-driven evaluation. Once a submission form is submitted, a member of the community with access to PRIME-RE’s backend can opt to include it on the website. It is up to the community to evaluate a resource’s usefulness. A simple community support rating system has been implemented to allow anyone to express support for a resource by ‘liking’ it. This system could help the users of PRIME-RE to navigate the collection of resources and find tools that have accumulated a lot of community support. If concerns arise, they can be discussed through any of the offered communication channels. Direct community discussions are available through a dedicated “prime-re” chat-channel on the Brainhack Mattermost server (https://mattermost.brainhack.org/brainhack/channels/prime-re), and more conventional mailing-list or forum style discussion is available using the Neurostars forum (https://neurostars.org) with the tag “prime-re“. The Neurostars forum is hosted by the International Neuroinformatics Coordinating Facility (INCF) and already has a broad user-base of neuroscience researchers. It is also possible to submit a contact form with comments, questions, or suggestions directly to PRIME-RE’s backend, and often also to individual developers of a resource through their provided communication information. To facilitate collaboration among its users, PRIME-RE also partners with the BrainWeb initiative (https://brain-web.github.io).

At the moment, PRIME-RE has two main components (Fig 1A), 1) the primary website with categorized resource listings, and 2) an evolving wiki to document the challenges and best practices of NHP imaging. The resource categories that are currently listed on the PRIME-RE website are: 1) Templates and atlases; 2) General analysis; 3) Structural analysis; 4) Functional analysis; 5) Diffusion analysis; 6) Data sharing; and 7) Software packages. The results section of this paper will present an overview of some of the custom tools listed under these categories at the time of writing. The Software Packages section contains a list of commonly used neuroimaging packages that were not specifically developed for NHP neuroimaging. Many of the specialized tools integrate with some of these packages or employ some of their subfunctions. The main website also provides links to the above-mentioned communication channels and to the wiki. The wiki contains a primer on NHP neuroimaging that documents common challenges and potential solutions. It has its own version controlled back-end and is currently open for contributions for anyone with a GitHub account. While the current wiki already constitutes a team effort, the document is presented as a starting point that could potentially evolve into a ‘best practices’ guide with dynamic involvement of the broader NHP neuroimaging community.

**Figure 1.**
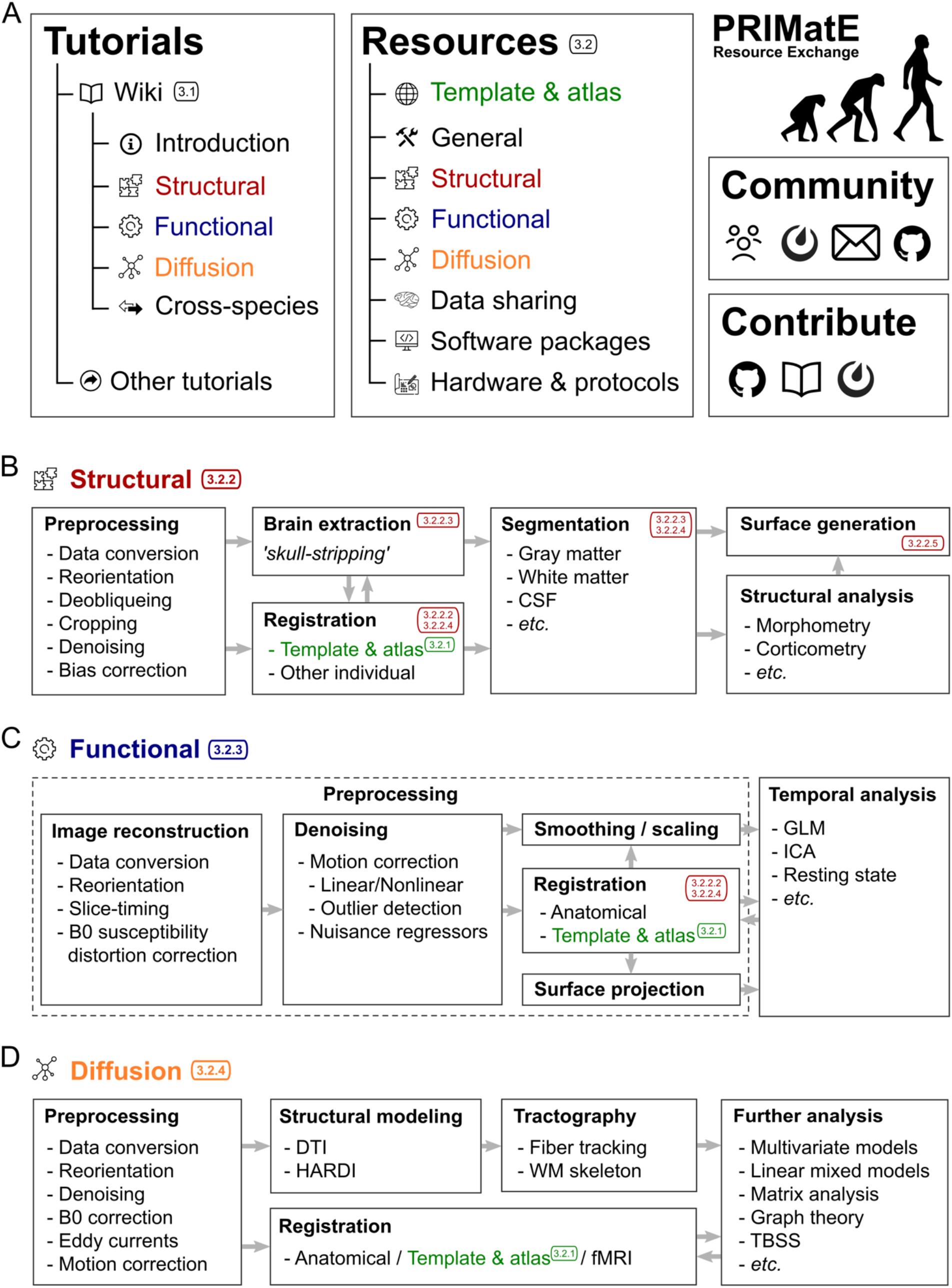
Organization of PRIME-RE and common NHP-neuroimaging analysis workflows. **A)** The main components of the PRIME-RE website (https://prime-re.github.io) are the Tutorials and Resources sections. The Tutorials section has a community-driven Wiki explaining the basics of NHP neuroimaging as well as links to external tutorials. The Resources section provides descriptions of the various tools, with links to the relevant files, documentation, and manuscripts. Both the wiki and the Resources section are organized thematically (e.g., structural, functional, diffusion), shown here in different colors. **B-D**) Example workflows for the analysis of structural (B), functional (C), and diffusion (D) NHP neuroimaging data, indicating common analysis steps. The colored numbers refer to section numbers in this paper that describe tools and pipelines developed to address these analysis steps. Note that there is not a standardized workflow for these types of analyses and that individual researchers may include or omit certain steps and proceed in an order that is fitting for their own needs and circumstances. The website also includes a Community section, where NHP neuroimaging researchers and developers can connect and find relevant literature, and a Contribute section that explains how people can get involved in the PRIME-RE initiative through GitHub, the Wiki, or the Mattermost channel.

Most of the tools that are currently listed on PRIME-RE were developed for macaque neuroimaging, but the platform aims to be an inclusive place where resources for all non-human primate species can be found. Several tools for marmoset neuroimaging are already listed, but the species diversity is likely to further increase as PRIME-RE evolves.

## 3. Results

The first version of the PRIME-RE platform went live in November 2019. It currently describes over 30 custom resources from contributors all over the world and houses an extensive primer on NHP neuroimaging in the form of a wiki with references to the resources listed on PRIME-RE. It also provides a comprehensive overview of neuroimaging tools that are not necessarily tailored towards NHP neuroimaging (although some of these packages do offer built-in solutions to facilitate working with non-human data). Here, we will present a synopsis of the wiki document (https://github.com/PRIME-RE/prime-re.github.io/wiki) and an overview of some of the specialized resources that are part of PRIME-RE at this time.

### 3.1 The wiki: A primer on NHP-MRI

The wiki-pages start with sections that address the various motivations for NHP neuroimaging and some of the common challenges encountered in data collection (Farivar and Vanduffel, 2014). These sections include study preparation issues, such as obtaining ethical approval for your study, responsible animal housing, handling, surgery, and transportation. The choice of coil type is discussed and the consequences of using custom coils are clearly explained. Other discussed challenges in data acquisition include non-standard body orientations, subject motion, the use of anesthesia or contrast agents, variable fields-of-view, and a reduced signal-to-noise-ratio (SNR) as a result of having to scan small NHP brains at higher resolution than human brains at the same magnetic field strength.

The next section briefly addresses the topic of data organization. Standardization efforts in human neuroimaging have yielded the Brain Imaging Data Structure (BIDS) standard, a widely adopted file-naming and data-structure convention that facilitates data and resource sharing. Whereas a range of tools on PRIME-RE understand data in BIDS format, the compatibility of NHP data with the BIDS structure is not perfect, again due to some unique challenges in NHP neuroimaging that are not present in human neuroimaging. There are, however, ongoing efforts to either expand the BIDS format so that it can better incorporate the idiosyncrasies of NHP data, or create a derivative NHP version that fulfills specific requirements to this research field.

The remainder of the wiki largely follows the categories that are also used in PRIME-RE’s resource listings. There are separate sections for structural, functional, and diffusion data processing (Figure 1A). The section on structural data processing includes data processing steps, such as orientation correction, de-obliqueing, cropping, denoising and averaging multiple volumes or images. It lists several options to handle bias-correction, brain extraction (skull-stripping), and segmentation. It also links to several pipelines available from PRIME-RE that offer more or less complete solutions to a range of these issues.

The section on functional data processing points out that this step is in fact not that different from human neuroimaging since the same types of statistical models can be applied to different species. One major difference that does exist is the extent to which data need to be pre-processed. NHP data, especially data from awake NHP neuroimaging studies, tend to require a lot more motion and distortion correction than the typical human dataset. The section on diffusion-weighted imaging discusses the differences between *in vivo* and *post-mortem* (*ex vivo*) diffusion MRI and their consequences for data analysis. Several caveats and solutions for tractography in NHPs are also laid out.

The wiki ends with a section on cross-species comparison. While challenging, this is also one of the most powerful uses of NHP neuroimaging since it has the potential to yield important translational or evolutionary insights. The section explains the recently developed common feature space approach (Mars et al., 2018a, 2018b), as well as comparisons based on activity dynamics (Mantini et al., 2012), and brain matching based on homologous sulci (Auzias et al., 2011). Several resources, suitable for the processing of NHP data, are suggested to implement these approaches.

The PRIME-RE wiki is not meant as a definitive guide to NHP neuroimaging, but rather as a community curated and dynamically updated collection of ‘best’ (or ‘optimal’) practices. Improvements and additions from the community are highly encouraged.

### 3.2 PRIME-RE Resources

The resources that are currently listed on PRIME-RE are divided into a number of categories based on their general purpose (Figure 1). Below we highlight these categories and a selection of tools from each. Please consult the website for a complete and up-to-date overview of the available resources (https://prime-re.github.io/resources). All resources listed on PRIME-RE, including those described here, were developed and assessed with datasets collected in accordance with locally approved guides and legislation concerning animal wellbeing, including the NIH Guide for Care and Use of Laboratory Animals, the U.K. Animals (Scientific Procedures) Act, 1986, European Directive 2010/63/EU, and the Australian Code for the Care and Use of Animals for Scientific Purposes.

#### 3.2.1 Templates & Atlases

Alignment of an individual’s (f)MRI data to a standard template space provides several advantages. Firstly, it provides a detailed anatomical image for evaluating and presenting functional results or other maps. Secondly, it presents a standard orientation and coordinate system for reporting findings, comparing results across subjects and labs, and aligning data for group analysis. Thirdly, templates frequently include ancillary datasets such as brainmasks, tissue segmentation masks, anatomical atlases, surfaces, morphometry, and data from other modalities that put a variety of analysis tools at the user’s disposal. These resources can eliminate time-consuming manual processing and encourage the use of consistent and reproducible processing streams. For example, a brainmask can be used for template-based brain extraction in the subject’s space or an atlas can be used for a region of interest (ROI) analysis in the template or subject space.

The choice and adoption of a standard template space is becoming especially relevant now that data sharing initiatives make it much more feasible to obtain larger sample sizes. For human (f)MRI, the MNI152 template (Fonov et al., 2011) provides a widely adopted standard volumetric space, while the fsaverage (Freesurfer average) (Fischl, 2012) template is a popular standard space for surface-based analysis. For non-human primate (NHP) MRI, there are a variety of templates. PRIME-RE currently lists several macaque and marmoset templates, as well as a mouse lemur template. In addition to species, templates differ with regard to imaging modality, *in vivo* versus *ex vivo* scanning, single subject versus population average, field strength, resolution, etc. For the rhesus macaque, the NIMH Macaque Template (NMT) has been widely adopted and has been incorporated in a number of the resources on PRIME-RE (Seidlitz et al., 2018). An updated version of this *in vivo* population template (NMT v2) is presented in this special issue and described below (Jung et al., this issue). The NMT v2 and its associated average surface (see Section 3.2.2.5.1), together with their tissue segmentations and atlases, constitute a complete set of complementary volumetric and surface spaces for representing macaque data.

Two novel macaque atlases are presented in this special issue and detailed below. Both are defined on the NMT v2 and manually refined to this template. The first is an atlas of the cortex (CHARM, Jung et al., this issue) and the second is an atlas of the subcortex, covering the forebrain, midbrain, and hindbrain (Hartig et al., this issue). These complementary atlases are arranged hierarchically, describing regions at various levels of granularity. Though these atlases have been tailored to the NMT v2, they can be aligned to previous macaque templates already used by some labs, and atlases in those templates can similarly be aligned to the NMT v2. The RheMAP resource, which is described in the subsequent section on Structural MRI Tools (see Section 3.2.2.2.2), stores nonlinear alignment warps between various macaque anatomical templates to facilitate such conversions between template spaces (Klink and Sirmpilatze, 2020; Sirmpilatze and Klink, 2020).

Two marmoset resources are also included in PRIME-RE. The Marmoset Brain Mapping Project is a multimodal high-resolution template and atlas of the marmoset brain that includes white matter maps based on diffusion MRI data. The most recent version is a population template based on *in vivo* scans that includes surfaces (Liu et al., this issue). In addition, the Marmoset Brain Connectivity Atlas (Majka et al., this issue) provides a large collection of anatomical tracer data and histological material in an interactive platform, equipped with various analysis tools.

##### 3.2.1.1. NIMH Macaque Template (NMT v2) and Hierarchical Atlas of the Cortex (CHARM)

*A macaque template in stereotaxic coordinates with a multi-scale cortical atlas.*

The National Institute of Mental Health (NIMH) Macaque Template (NMT) is an anatomical MRI template of the macaque brain that serves as a standardized space for macaque neuroimaging analysis (Seidlitz et al., 2018). The recently released NMT version 2 (Jung et al., this issue) provides a complete overhaul of the NMT, including a fully-symmetric template in stereotaxic orientation (i.e., in the Horsley-Clarke plane) (Horsley and Clarke, 1908). Coordinates in this stereotaxic space are measured from ear bar zero (i.e., the intersection of the midsagittal plane and a line through the interaural meatus). The adoption of stereotaxic orientation and coordinates will assist users with surgical planning and facilitates reporting of coordinates commensurate with those used with other techniques (e.g. electrophysiology, intracerebral injection).

The NMT was created by iteratively registering T1-weighted scans of 31 adult rhesus macaque brains to a working template, averaging the nonlinearly registered scans, and then applying the inverse transformations to bring the working template closer to the group average (Seidlitz et al., 2018). The symmetric NMT was generated through the same process except that each subject’s anatomical was input twice, once in its true orientation and once mirrored about the midline. Modifications to the scan averaging and postprocessing have improved template contrast and spatial resolution. Brain masks, segmentations, and various other tissue masks are provided (Figure 2A-C). The availability of symmetric and asymmetric variants of the NMT v2 allows users to choose the version that best suits their analysis. Other template variants include an expanded full head field-of-view and a lower resolution version for faster processing (0.5 mm isotropic, down-sampled from 0.25 mm isotropic). Surfaces for NMT v2, generated using the new CIVET-Macaque platform (Lepage et al., this issue), are provided for easy data visualization. For surface-based group analysis, the NMT average surface, which comes with anatomical labels from the D99 and CHARM atlases, can serve as a surface-based registration target for representing cortical surfaces in a common framework regardless of the processing pipeline.

**Figure 2.**
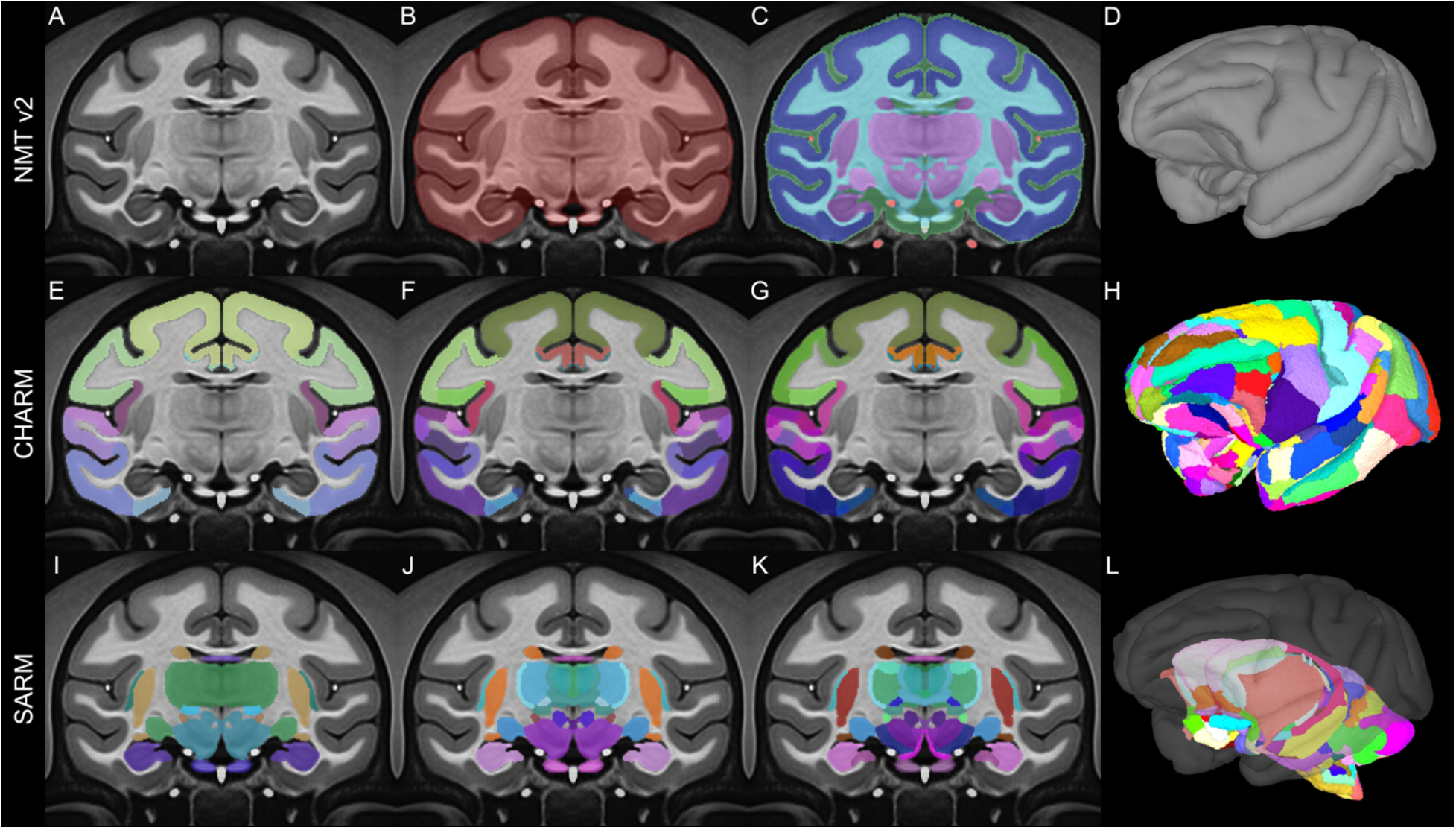
Macaque Templates and Atlases. The symmetric variant of the NIMH Macaque Template v2 (NMT v2), with tissue segmentation (top row) and anatomical regions (middle and bottom rows) depicted in color on a coronal slice (11 mm anterior to ear bar zero) and the pial surface (right column). The NMT v2 includes **A**) the population average volume; **B**) a manually refined brain mask that indicates which voxels contain brain tissue; **C**) a 5-class tissue segmentation that differentiates cerebrospinal fluid (green), gray matter (dark blue), white matter (light blue), subcortex (purple), and vasculature (red); and **D**) rendering of surfaces. The middle row shows the Cortical Hierarchy Atlas of the Rhesus Macaque (CHARM) and the bottom row shows the Subcortical Atlas of the Rhesus Macaque (SARM). The atlases provide anatomical labeling of cortical and subcortical regions, respectively, at six progressively finer spatial scales. Parcellations for **E,I**) Level 2, **F,J**) Level 4, and **G,K**) Level 6 are shown. **H,L**) Surface representation of the finest (Level 6) parcellations.

Atlases are an important aspect of group and ROI-based analyses. The NMT v2 package comes with multiple anatomical atlases that have been manually tailored to the template’s morphology. These atlases include the D99 atlas (Reveley et al., 2017) and the new Cortical Hierarchy Atlas of the Rhesus Macaque (CHARM; Jung et al., this issue). The latter is a novel anatomical parcellation of the macaque cerebral cortex, where the cortical sheet is subdivided into six-levels of increasingly fine-grained parcellation (Figure 2D-F). The broadest level consists of the four cortical lobes and the finest level is based on the D99 atlas, with modifications that make the regions more robust when applied to low resolution (e.g. fMRI) data. Different scales of CHARM can be combined so that, for example, a tracer injection or the seed region for a resting state analysis can be described using a fine scale, while the region’s anatomical or functional connectivity can be succinctly described on a broader scale. This way, whole brain data can be characterized on a spatial scale that is most suitable to a study’s findings. Users can also select a CHARM level *a priori* based on how many regions it contains, and thus control the required degree of multiple comparison correction for their analysis.

While the NMT v2 works with any neuroimaging platform that accepts NifTI/GifTI format, the template has been designed to have additional functionality within the AFNI ecosystem (Cox, 1996). Enhancements include support for the *NMT2* standardized space, recognition of the NMT v2 in the @animal_warper alignment pipeline (see section 3.2.2.4.2) and the functional processing pipeline generator afni_proc.py (see section 3.2.3.1), and downloadable demos showing how AFNI can perform structural and functional analyses (task-based or resting state) using the NMT v2. See the accompanying article by Jung et al. (Jung et al., this issue) for further information.

##### 3.2.1.2. Subcortical Atlas of the Rhesus Macaque (SARM)

*A complete MRI atlas of the macaque subcortex suited for neuroimaging.*

The Subcortical Atlas of the Rhesus Macaque (SARM) is an anatomical parcellation of the entire subcortex tailored for magnetic resonance imaging (MRI) (Hartig et al., this issue). The regions-of-interest (ROIs) are hierarchically organized, with grouping levels suited for both fine structural and spatially broader functional analyses. SARM aims to facilitate the identification, localization, and study of neural interactions involving subcortical regions of the brain.

A high-resolution structural MRI of an *ex vivo* rhesus monkey brain was segmented into 180 anatomical ROIs by matching distinctly contrasted structures with their histological counterparts. ROIs followed the recently updated nomenclature and delineations of the Rhesus Monkey Brain in Stereotaxic Coordinates atlas (Paxinos et al., in preparation). The *ex vivo* MRI was nonlinearly registered to the symmetric population-level template of the rhesus macaque NMT v2 (see section 3.2.1.1 and (Jung et al., this issue). The SARM was warped using Advanced Normalization Tools (ANTs; (Avants et al., 2008) and refined using AFNI (Cox, 1996). ROIs were delimited using the tissue segmentation masks for the NMT v2, smoothed by computing the mode over neighboring voxels, and individually refined by hand to match the average structural template. The SARM is organized into 6 hierarchical levels. The finest level presents each individual ROI (Figure 2G), whereas the broadest level corresponds to developmental subdivisions (i.e., tel-, di-, mes-, met-, and myel-encephalon). The SARM was validated in 3 monkeys with a functional MRI (fMRI) paradigm known to activate the lateral geniculate nucleus (LGN) (Logothetis et al., 1999).

The SARM offers neuroimaging researchers a complete subcortical segmentation of the rhesus macaque brain. The atlas can be used for a range of structural and functional analyses and has been specifically tailored to work with the CHARM and NMT v2 (Jung et al., this issue). The SARM is also available to download via Zenodo, and has conveniently been incorporated with the atlases available in AFNI. Additional information is available in the accompanying article (Hartig et al., this issue).

##### 3.2.1.3. Marmoset Brain Mapping Project (MBM)

*MRI-based marmoset brain atlases and tools for neuroimaging and connectome studies*

The Marmoset Brain Mapping Project (www.marmosetbrainmapping.org) includes three atlases and templates:

1)Version 1 focuses on cortical parcellations (Liu et al., 2018). A 3D digital gray matter atlas was constructed from high-resolution (150 μm isotropic) *ex vivo* MRI images, including magnetization transfer ratio (a T1-like contrast), T2w images, and multi-shell diffusion MRI (dMRI). Based on the multi-modal MRI contrasts, 54 cortical areas and 16 subcortical regions were manually delineated on one hemisphere of the brain using a data-driven approach that was developed to minimize manual drawing errors. The 54 cortical areas were then merged into 13 larger cortical regions according to their locations to yield a coarse version of the atlas, and also subdivided into 106 regions using a dMRI connectivity-based parcellation method. A Paxinos-style cortical atlas (Majka et al., 2016) was fused and refined into the high-resolution MRI template to provide interoperability to other marmoset databases. The atlas set provides a readily usable multi-modal template space with multi-level anatomical labels that can facilitate various neuroimaging studies of marmosets.

2)Version 2 focuses on fine-detailed white matter pathways (Liu et al., 2020). In this version, *ex-vivo* MRI data of the marmoset brain was collected with the highest spatial resolution available to date, including 80 μm and 64 μm multi-shell dMRI, 80 μm MTR, and 50 μm T2w and T2*w images. The data allowed building a fine-grained 3D white matter atlas, which depicts many fiber pathways that were either omitted or incorrectly described in previous MRI datasets or atlases of the primate brain. By combining dMRI tractography and neuronal tracing data (Majka et al., 2020), a detailed fiber pathway mapping of cortical connections is provided. The white matter atlas, fiber pathways maps and dMRI data, including both raw and processed data, are publicly available on the project’s website (Figure 3A,B).

**Figure 3.**
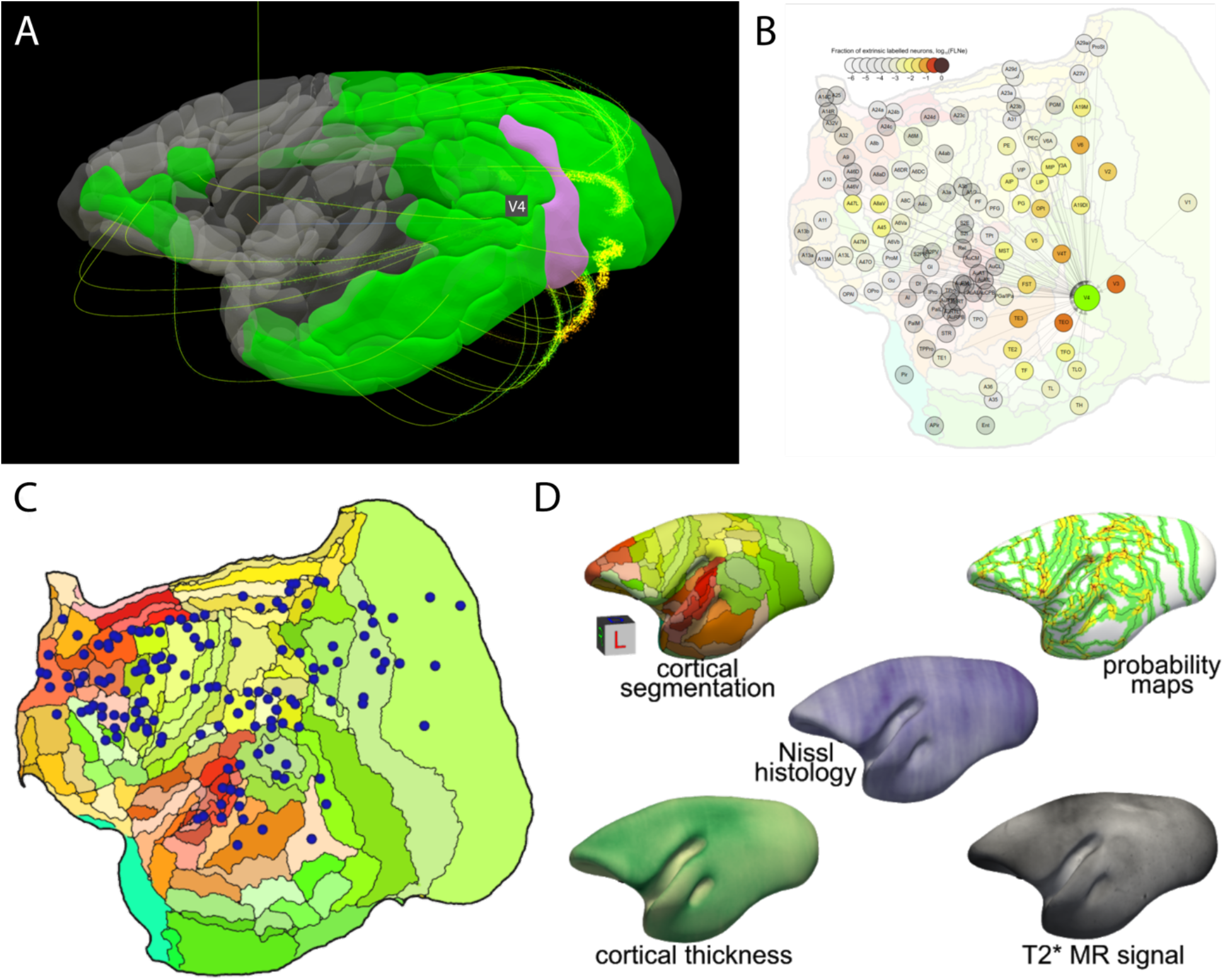
Marmoset atlases. **A-B)** Example visualizations available on the website of the Marmoset Brain Mapping Project. **A**) 3D-viewer that demonstrates the connectivity profile of an area selected by clicking (here V4). **B**) Comparison with the weighted and directed connectivity graph based on the results of monosynaptic retrograde fluorescent tracer injections from the Marmoset Brain Connectivity Atlas (https://analytics.marmosetbrain.org/graph.html). The graph view highlights the spatial relations between connected areas. Clicking a node reveals further information, including an average connectivity profile, interactive visualizations of the data for each injection, and metadata. **C-D)** Marmoset Connectivity Atlas**. C**) Locations of 143 tracer injections registered to a stereotaxic template illustrated in a 2-dimensional flat-map of the marmoset cortex (C). **D**) The main data layers of the histology-based average morphology of the adult marmoset brain showcasing the interoperability of this dataset with neuroimaging research (T2* MR signal).

3)Version 3 focuses on population standard templates and cortical surfaces (Liu et al., this issue). While versions 1 and 2 were based on a few *ex vivo* brain samples, and lacked essential functionalities for *in vivo* studies of large animal cohorts, version 3 is based on *in vivo* population data. Standard templates are derived from multi-modal data of 27 marmosets, including multiple types of T1w and T2w contrast images, DTI contrasts, and large field-of-view MRI and CT images. Multi-atlas labeling of anatomical structures was performed on the new templates and highly accurate tissue-type segmentation maps were constructed to facilitate volumetric studies. Fully featured brain surfaces and cortical flat maps facilitate 3D visualization and surface-based analyses with most surface analyzing tools. The population-based template will significantly aid a wide range of neuroimaging and connectome studies that involve across-subject analysis.

##### 3.2.1.4. Marmoset Brain Connectivity Atlas

*Marmoset anatomical tracer data and connectivity analysis tools integrated with an atlas and histological material.*

The Marmoset Brain Connectivity Atlas (Majka et al., 2020) allows exploration of a growing collection of data from retrograde tracer injections in the marmoset neocortex. At present, it includes data from over 140 experiments covering almost 50% of currently recognized areas of marmoset cortex, encompassing subdivisions of prefrontal, premotor, superior temporal, parietal, retrosplenial and occipital complexes. Data from different animals are presented against high-resolution images of the underlying histology, and registered to a template based on the Paxinos et al. (Paxinos et al., 2012) stereotaxic atlas using an algorithm guided by an expert delineation of histological borders. The portal (http://www.marmosetbrain.org) implements best practices in terms of sharing the connectivity data. The resource provides access to primary experimental results, as well as connectivity patterns that are quantified according to the currently accepted parcellation of the marmoset cortex. To enable graph-based network analyses, the portal incorporates tools for data exploration relative to cytoarchitectural areas (http://analysis.marmosetbrain.org), including statistical properties such as the fraction of labeled neurons, the percentage of supragranular neurons, and geodesic distances between areas across the white matter. Importantly, it also provides the cellular connectivity data in a purely spatial (parcellation-free) format, including the stereotaxic coordinates of ~2 million neurons labeled by the tracer injections. These results can be downloaded in 3D volume format in the template space (Majka et al., this issue), so they can be compared with MRI-based topologies. The portal also makes data available in a machine-mineable way (http://analytics.marmosetbrain.org/wiki/api) to facilitate large-scale models and simulations.

Apart from the tracer-based connectivity data (Figure 3C), the portal hosts a collection of other unique datasets typically inaccessible in non-invasive imaging, such as high-resolution images of cytoarchitectural characteristics for each of the marmoset cortical areas, stained with a neuron-specific antibody (Atapour et al., 2019) (http://www.marmosetbrain.org/cell_density). Moreover, a unique histology-based average morphology of the adult marmoset brain provides the basis for probabilistic registration of digital datasets to cytoarchitectural areas (Majka et al. this issue; http://www.marmosetbrain.org/nencki_monash_template). Spatial transformations to other marmoset brain templates are included to enable prompt integration with magnetic resonance imaging (MRI) and tracer-based connectivity studies (Figure 3D).

#### 3.2.2. Structural MRI Tools

##### 3.2.2.1. Overview

Pre-processing of structural and functional MRI involves alignment of volumetric data across scanning sessions or with a species-specific anatomical template/atlas in a standardized space; segmentation (i.e., differentiation) of the brain from skull, muscle, and other tissues (a.k.a. “skull-stripping”); segmentation of different tissue classes within the brain (e.g. gray and white matter). Additionally, volumetric cortical data are sometimes represented on a flat map or surface map by creating individualized surfaces or projecting data to a common surface. Here we briefly describe some PRIME-RE resources for handling volumetric data (Figure 1B). Typically, anatomical scans are analyzed in detail and the results are then also applied to any functional data collected in the same scanning session.

Some of the structural tools described perform a single pre-processing step, allowing them to be flexibly used with other tools. Examples of dedicated alignment tools are Reorient, which does rigid body alignment and cropping the borders of a scan, and RheMAP, which provides the non-linear warps between various macaque templates. Dedicated segmentation tools include the interactive Thresholdmann and the machine-learning based UNet, both of which generate binary brain masks, and BrainBox, which is well suited for the collaborative development of binary or multi-valued masks.

Other resources perform multiple steps within a single software environment or by calling on various other tools. For example, Macapype uses several tools interchangeably to perform alignment, normalization, and segmentation operations. Similarly, AFNI’s @animal_warper computes and applies a nonlinear alignment between a subject and a template and can warp the template’s masks to create a brain mask, tissue segmentation, and atlas parcellation for the individual. For other tools, alignment and segmentation are precursors to additional steps such as surface generation (e.g., CIVET-Macaque, PREEMACS) or functional analysis (e.g., C-PAC).

PRIME-RE offers four tools for generating surfaces from volumetric monkey data. The NHP-Freesurfer, PREEMACS, and Precon_all tools all utilize the Freesurfer software package in conjunction with several other tools. In contrast, CIVET-macaque is a novel adaptation of CIVET for use with monkey data. It can be used to characterize surface morphology such as cortical thickness (Lepage et al., this issue). All four programs require a T1-weighted anatomical scan. Both CIVET-macaque and PREEMACS can take an optional T2-weighted scan as well, and both provide quality control tools.

##### 3.2.2.2. Alignment and registration tools

Having properly oriented MRI datasets is important for many neuroimaging workflows and often crucial for further segmentation or registration across functional, anatomical and template images. For non-human scans (both *in vivo* and *ex vivo*), orientations and field-of-views can vary tremendously, which means that a correction through reorientation and cropping is often required before any standard tools can be used. Group analysis of (f)MRI data is commonly performed in a standard template space to which individual data are registered. Here we describe tools for aligning datasets to a template or to each other as well as precalculated registrations between templates (RheMAP). These tools include Reorient, which aids in visual rigid body alignment, and nonlinear alignment tools such as macapype and @animal_warper (see Section 3.2.2.4).

###### 3.2.2.2.1. Reorient

*An intuitive web tool for reorienting and cropping MRI data.*

Reorient (https://neuroanatomy.github.io/reorient) is an open source web application for the intuitive manual alignment and cropping of MRI NifTI volumes. The MRI scan is dragged onto a web interface and visualized in an interactive stereotaxic viewer. Users can then translate and rotate the brain by simply dragging inside the three view planes. An adjustable selection box determines the cropping of an image. Resulting affine matrices, selection boxes, as well as the reoriented and cropped volume can be saved. The affine matrix and selection box can then be used in scripted workflows to make these steps reproducible. Existing rotation matrices can also be loaded or appended in the web interface. Reorient complements existing tools by providing an intuitive approach for manual image reorientation and all components for a fully reproducible workflow. It has been used extensively to reorient scans from many different vertebrate species, including 60 different primate species.

###### 3.2.2.2.2. RheMAP

*Precomputed nonlinear warps between common rhesus macaque brain templates.*

The generation of accurate nonlinear registrations between different brain images is a time-consuming and computationally expensive operation. The RheMAP project was created to generate a set of pre-computed nonlinear registration warps that allow the direct remapping of (f)MRI data across different common macaque template brains. The non-linear warps were generated using ANTs (Avants et al., 2008) and the Python code that was used in RheMAP to compute the warps is available on GitHub (https://github.com/PRIME-RE/RheMAP) together with documentation and example code that explains how to use the warps to remap data between template spaces or register a single anatomical scan to one or more template spaces (Klink and Sirmpilatze, 2020). The dataset of computed warps is freely available for download from Zenodo (Sirmpilatze and Klink, 2020). Registration quality can be visually assessed with the help of images that are distributed with the code and the dataset. RheMAP already includes a comprehensive set of templates — NMT (Jung et al., this issue; Seidlitz et al., 2018), D99 (Reveley et al., 2017), INIA19 (Rohlfing et al., 2012), MNI macaque (Frey et al., 2011) and Yerkes19 (Donahue et al., 2016) — but the provided code allows for easy inclusion of additional template brains as well. Moreover, the general warp computation workflow can be easily adapted for other animal species, since the underlying ANTs registration functions do not rely on prior knowledge about brain size and shape.

##### 3.2.2.3. Segmentation tools

Obtaining appropriate tissue masks (brain vs. surrounding tissue, or different tissue types within the brain itself) can be particularly difficult in non-human brain imaging, as standard automatic segmentation tools struggle with surrounding muscle tissue, the skull, and the strong intensity gradients that are often present. PRIME-RE lists several tools that can facilitate segmentation of NHP brain scans.

###### 3.2.2.3.1. Thresholdmann

*A web tool for interactively creating adaptive thresholds to segment NifTI images.*

A simple intensity threshold can often provide a good initial guess for tissue segmentation, but in the presence of strong intensity gradients, a threshold that works well for one brain region can easily fail elsewhere in the same brain image. Thresholdmann (https://neuroanatomy.github.io/thresholdmann) is an open source web tool for the interactive application of space-varying thresholds to NifTI volumes. It does not require any software downloads or installation and all processing is done on the user’s computer. NifTI volumes are dragged onto a web interface where they become available for visual exploration in a stereotaxic viewer (Figure 4A). A space-varying threshold is then created by setting control points, each with their own local threshold value. The viewer is initialized with one control point at the center of the brain. The addition of further control points produces a space-varying threshold obtained through radial basis function interpolation. Each local threshold can be adjusted in real time using sliders. Finally, the thresholded mask, the space varying threshold and the list of control points can be saved for later use in scripted workflows. Thresholdmann complements a variety of existing brain segmentation tools with an easy interface to manually control the segmentation on a local scale. The resulting masks can serve as starting points for more detailed manual editing using tools such as BrainBox (https://brainbox.pasteur.fr) or ITK Snap (http://www.itksnap.org). The interactive approach is especially valuable for non-human brain imaging data, where it has been successfully used to create initial brain masks for a variety of vertebrate brains – including many NHP datasets (Heuer et al., 2019) – as well as developmental data.

**Figure 4.**
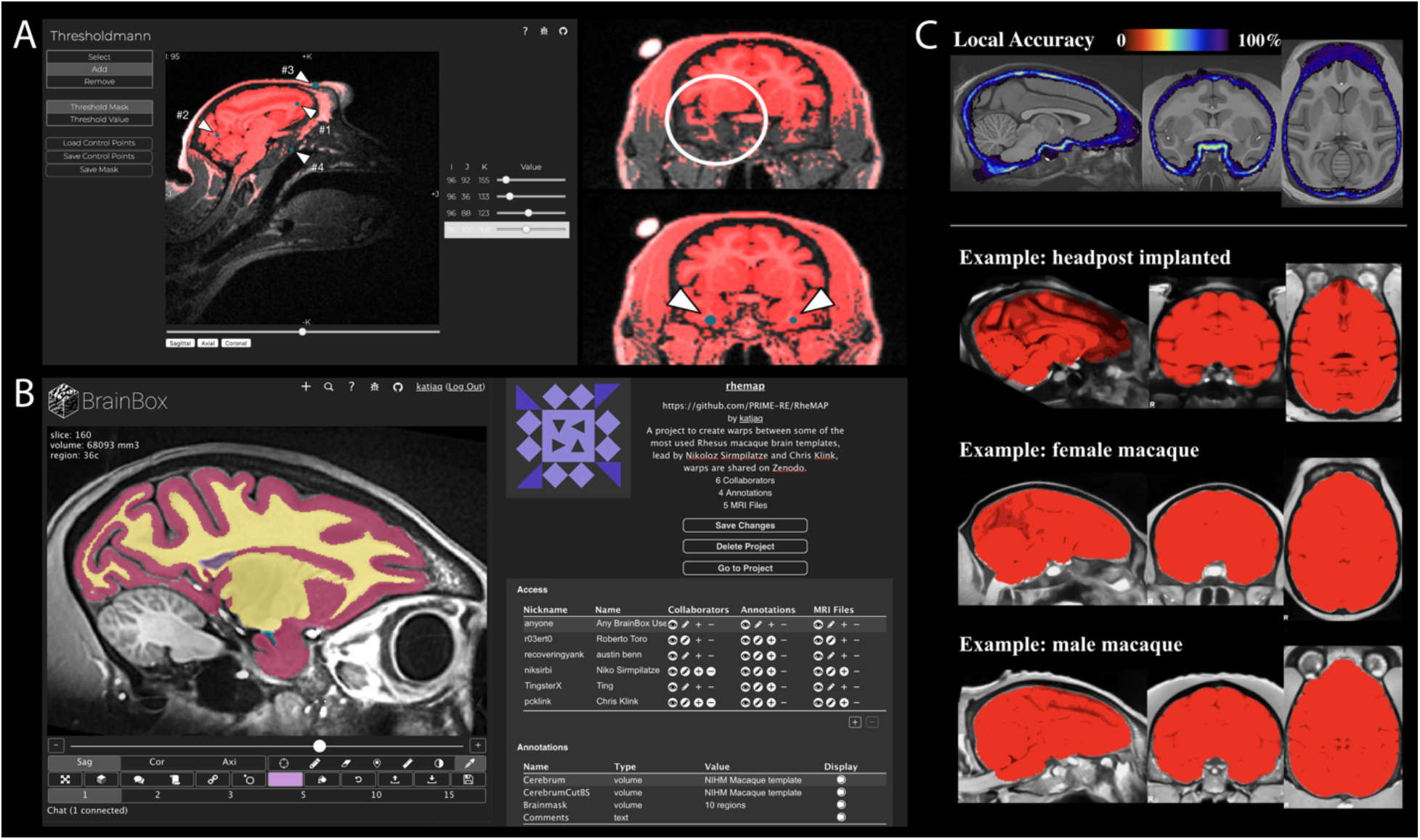
Segmentation tools. **A**) Thresholdmann is a brain extraction tool that creates a mask from those voxels that are brighter than a spatially varying threshold. The threshold is constrained by control points (blue dots flagged with number arrows) that are added by clicking at locations in the web viewer (left). Adjustable sliders to the right control the threshold in the vicinity of each control point. In the examples shown to the right, the mask initially (top) excludes ventral brain areas but is filled out (bottom) by the addition of ventral control points. **B**) Brainbox is a web-based tool for collaborative brain segmentation. In the interface, each MRI (left) has a page where information on all projects utilizing this dataset is centralized. Under project settings (right), users can assemble datasets into projects, add collaborators, manage access rights, and add annotation layers. **C**) The UNet tool uses a neural network model approach for brain extraction. Local accuracy at each voxel is estimated by the regional Dice coefficient (top). Bottom panels show results for three example subjects from the PRIME-DE repository, with regions assessed to be brain tissue shown in red.

###### 3.2.2.3.2. BrainBox

*A web application for real time, collaborative visualization, annotation & segmentation*

Automatic brain segmentations often need to be manually adjusted and in some cases a complete manual segmentation is required. Manual segmentation can however be difficult, time consuming, and it is often performed by researchers working in isolation, even for shared public data. BrainBox (https://brainbox.pasteur.fr) is an open source tool for the visualization, collaborative curation and analysis of neuroimaging data. BrainBox makes it possible for several researchers to simultaneously work on the same dataset in a shared virtual working space. BrainBox can be used to visualize and provide a layer of human annotation to any dataset available on the Web, for example, to data stored in Zenodo, FigShare, Google Drive, Dropbox, Amazon S3, GitHub, or the Open Science Framework. After providing the URL of a dataset, BrainBox presents an interactive viewer, and a series of editing tools, e.g. for segmenting brain regions, erasing, or building 3-D models (Figure 4B). Several datasets can be combined into “projects” and project creators can invite collaborators. Access to the data can range from completely private to completely open projects. Finally, BrainBox implements a RESTful API (Application Program Interface) which allows programmatic access to all its data. For example, annotations and segmentations can be queried by a local python script, or even a shared notebook running on Google Colab.

BrainBox adds a layer of collaborative curation and analysis to shared neuroimaging datasets using only a web browser, allowing users to incrementally improve each other’s work. This increases scientific efficiency, improves public data quality, and reduces redundant effort. All PRIME-DE datasets (Milham et al., 2018) are already indexed in BrainBox projects, and ready for collaboration.

###### 3.2.2.3.3. UNet NHP brain extraction tool

*A convolutional neural network model for skull-stripping of NHP images.*

The UNet brain extraction tool provides a flexible brain labeling solution to facilitate the preprocessing of NHP neuroimaging data, particularly for multi-site datasets with different scan acquisition, quality, and surgical situations (e.g., with head-holder implants) (Figure 4C). The UNet model was initially trained on a large sample of human neuroimaging data and transferred to the NHP with additional macaque training samples from the PRIME-DE data repository (Milham et al., 2018). In order to run the tool with the default model, a user only needs to specify the input T1-weighted image. The full process requires ~20s on a GPU (NVIDIA GTX 1070TI, 700MB graphics card memory) and ~2-15 min on a CPU, which means it can easily be run on a personal computer. The current default model has been successfully applied to 136 macaque monkey scans from 19 sites on PRIME-DE with good performance (https://github.com/HumanBrainED/NHP-BrainExtraction). The tool also provides a module for the user to upgrade the UNet model for a specific dataset (e.g., raw T1w from a highfield scan, MP2RAGE, etc.). Unlike the traditional neuroimaging tools, no strong imaging background is required for the user to adjust processing parameters for a specific dataset. Instead, the tool only requires a small macaque image-set training sample (n=1-2) to upgrade the model and improve performance accuracy for that specific dataset.

##### 3.2.2.4. Alignment and Segmentation Pipelines

In addition to tools that specialize in a single step of structural analysis, there are also several pipelines listed on PRIME-RE that can take care of multiple structural analysis steps.

###### 3.2.2.4.1. Macapype

*An open multi-software framework for non-human primate anatomical MRI processing*

Macapype (https://github.com/Macatools/macapype) provides an open-source framework to create customized NHP-specific MRI data processing pipelines based on Nipype (Gorgolewski et al., 2011). Nipype is a widely-adopted Python framework for human MRI data processing, which provides wrappers of various commands and functions from well-known neuroimaging software (e.g., AFNI, FSL, SPM12, ANTs). In Macapype, custom scripts specific to NHP data processing are also wrapped. These include a brain extraction tool (AtlasBREX; Lohmeier et al., 2019) and a script for computing registrations between a subject brain and the NMT macaque template (NMT_subject_align.csh; Seidlitz et al., 2018). Users can thus flexibly construct customized pipelines by putting together various processing modules (and parameters) that are optimal for their dataset.

Macapype consists of several configurable modules for: 1) Data preparation steps, such as image reorientation, deoblique-ing, cropping, and the registration and averaging of multiple images; 2) Anatomical preprocessing steps like bias-correction with ANTs N4BiasCorrection; Tustison et al., 2010) and/or T1w x T2w bias field correction (Rilling et al., 2012), denoising using the adaptive non local means filter (Manjón et al., 2010), brain extraction (e.g. with AtlasBREX), and brain segmentation (e.g. using AntsAtroposN4 or SPM Old Segment).

Two examples of modular anatomical pipelines are already implemented in the distributed version of Macapype. They are customized for the preprocessing, brain extraction and segmentation of macaque anatomical data, and have performed robust segmentations on different datasets. The first pipeline creates a high quality segmentation in native space for surface reconstruction. Briefly, the pipeline takes the T1w and T2w images as inputs, and first applies cropping, bias-correction (via T1w x T2w and N4 intensity bias correction), and denoising (adaptive non-local means filtering) to improve image quality. Next, brain extraction is performed using AtlasBREX. For segmentation, transformations between the subject skull-stripped image and the NMT macaque template are first computed using the NMT_subject_align.csh script (Seidlitz et al., 2018), and then applied to register the NMT tissue segmentations to the subject image. Finally, segmentation is performed using AntsAtroposN4.sh to segment the subject image in native space, with the template segmentation as priors. The second pipeline reaches the same goal with different packages. This pipeline provides an iterative sequence for normalization (source to template space) of T1w and T2w, and provides segmentation with both SPM12 (OldSegment) and FSL FAST.

A docker file is included in the package to allow users to get a fully embedded version of the pipelines working on any computer without previous installation of MRI processing software. Both pipelines are compatible with BIDS (Gorgolewski et al., 2016) formatted datasets. The two pipelines have been demonstrated to achieve robust skull-stripping and segmentation on anatomical data from different primate species: Pipeline 1 was tested on both macaque (Milham et al., 2018) and marmoset datasets, while Pipeline 2 was tested on both human and macaque datasets. Documentation and an extensive description of the segmentation results from the macaque and marmoset brain extraction and segmentation are available on Github (https://macatools.github.io/macapype).

###### 3.2.2.4.2. @animal_warper

*AFNI program that aligns structural data to a template.*

@animal_warper is a bidirectional alignment program made for animal neuroimaging. While human neuroimaging has typically required a standard template, many animal researchers may prefer to keep their data in the native acquisition space and move atlas regions and tissue segmentations into the native space of the subject. Alternatively, data can be transformed to a standard template space to allow voxelwise group analysis and have the advantage of a standardized coordinate reporting system.

The @animal_warper program proceeds in a series of alignment steps. First, the center of the input dataset is moved to match the center of the template. Then, an affine alignment uses a 12-parameter transformation to match translation, rotation, shear and scale. Finally, a nonlinear warp is computed to align structural details to the template. The inverse warps and inverse affine transformations are computed and applied for the reverse direction of template to native space. “Follower” datasets, like ROIs drawn in the native space, can also be transformed into a target space. Datasets and ROIs typically follow from native to template space; templates, atlases and segmentations follow from template to native space.

The widely varying kinds of data used in animal imaging require different cost functions for alignment to flexibly deal with the voxel resolution and with the kinds of tissue contrast in each imaging modality. Several features of @animal_warper address some of the idiosyncrasies of animal alignment. First, the user can specify a “feature size” that controls blurring and cost functions. For many macaque MRI datasets, a value of 0.5 mm, which roughly matches the apparent voxel resolution, is a useful feature size. Other species, such as mouse, marmoset, and rat, may require a feature size that accommodates the finer voxel resolution typically used with these animals. Animal brain sizes can vary dramatically from each other and from any particular template, so a “supersize” option allows for up to a 50% difference in size. The program applies a kind of spatial regularization for ROI and atlas regions that goes beyond the typical nearest neighbor interpolation; a modal smoothing is applied to all transformed ROIs. This kind of smoothing assigns to each voxel the most common voxel value in a user-specified radius around it.

@animal_warper is developed within the larger AFNI software ecosystem. The computed transformations serve as input to a general fMRI processing pipeline implemented with afni_proc.py (see section 3.2.3.1). Output datasets in native space are viewable in the AFNI GUI and marked with the appropriate space. Data that have been transformed to a standard space have all the functionality of the “whereami” atlas command line and GUI. While NifTI datasets are marked as being in a scanner, Talairach, MNI or other space, @animal_warper adds an identifying space to the data. Atlas regions are generated in the native space both as volumes (Figure 5A) and as individual surfaces in GifTI format. Every region can be shown by itself or with any or all other atlas regions along with a simple surface rendition of the template in the native space of the subject. Quality control reports are generated as simple png images.

**Figure 5.**
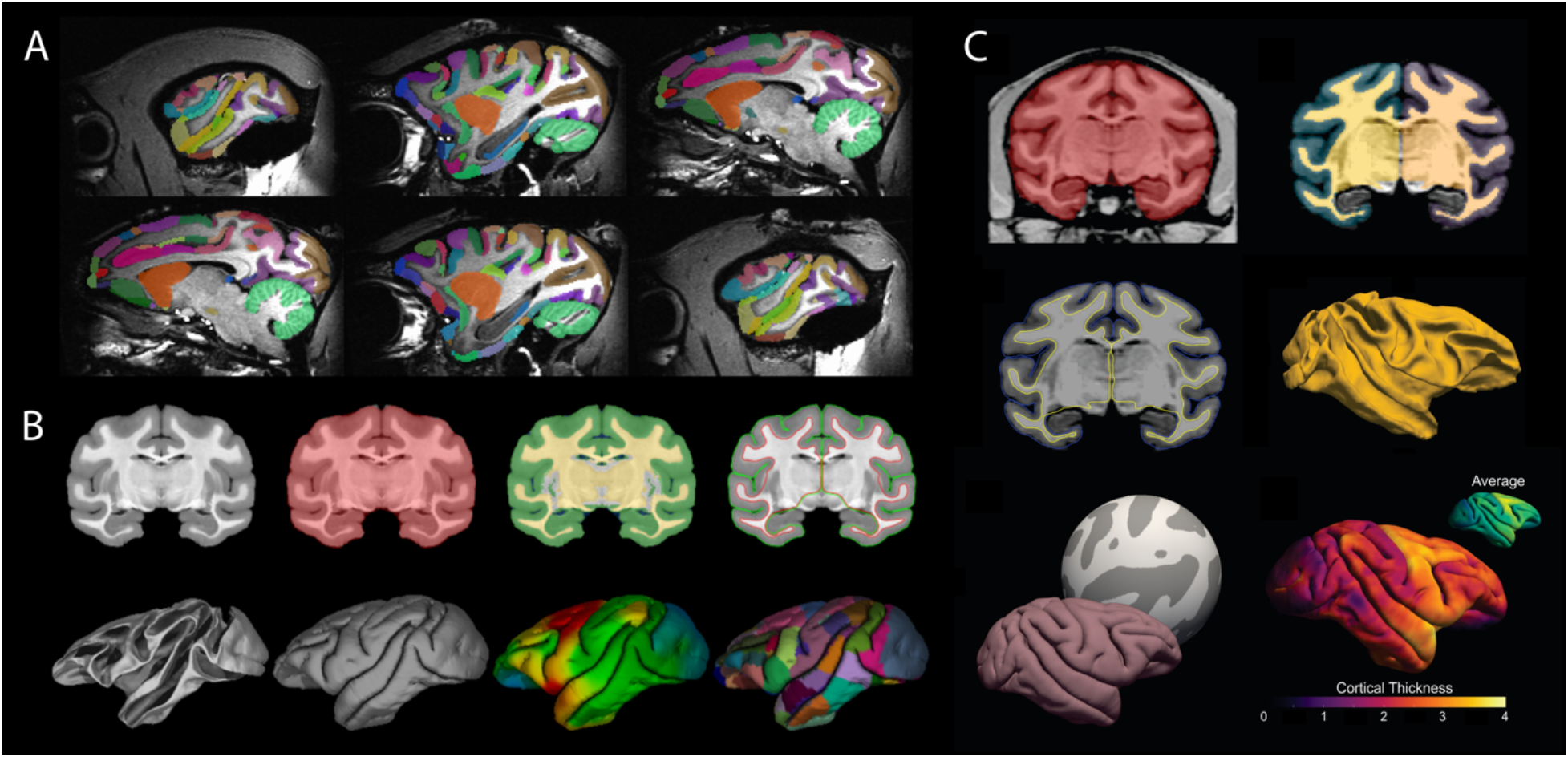
Anatomical MRI pipelines. **A**) Six sagittal slices illustrating the D99 atlas transformed to the native space of a macaque subject using @animal_warper. **B**) CIVET-macaque results for the D99 subject. Top row (left to right): anatomical scan, brain mask (red) from brain extraction, tissue classification (CSF in dark blue, cortical GM in green, portions of non-cortical GM in white, WM and other non-cortical GM in yellow) from segmentation, and generated cortical surfaces overlaid on the anatomical scan (WM surface in red, pial surface in green). Bottom row (left to right): 3D-renderings of the white and pial surfaces; cortical thickness indication from morphometric analysis (1.0 mm in light blue to 3.5 mm in red) and D99 surface parcellation (atlas) viewed on the pial surface. **C**) PREEMACS results. Top row: brainmask (red) created by a Deep Learning convolutional neural network model (left) and volumetric tissue segmentation (right). Middle row: white matter and pial surface estimation (left) and rendered white matter surface (right). Bottom row left: surface registration to the PREEMACS Rhesus parameterization template to obtain vertex correspondence (i.e., surface coregistration) between subjects. Bottom row right: individual cortical thickness estimation from morphometric analysis. The resulting surfaces can be analyzed in the geometric space of the PREEMACS average surface.

The program has been tested with a wide variety of the macaque data available on the PRIME-DE website (Milham et al., 2018). The AFNI website hosts the D99, NMT v1 (Seidlitz et al., 2018), the stereotaxic NMT v2 templates as well as the D99 atlas and the CHARM (see section 3.2.1.1 and Jung et al., this issue) and SARM (see section 3.2.1.2 and Hartig et al., this issue) hierarchical atlases.

##### 3.2.2.5. Surface generation pipelines

Surface visualizations offer a rich view of cortical fMRI activity. The generation of surface visualizations for non-human brains can however be rather involved. The pipelines below were specifically designed to facilitate this process.

###### 3.2.2.5.1. CIVET-macaque

*A pipeline for generating surfaces and cortical thickness maps from anatomical MRIs*

The CIVET pipeline for automated reconstruction of cortical surfaces from *in vivo* human MRI has been extended for processing macaque brains. CIVET-macaque requires only a T1-weighted MRI image to generate high resolution cortical surfaces, capable of intersubject surface-based coregistration with per-vertex correspondence between gray and white matter surfaces. Preprocessing and surface reconstruction were customized for macaques to address species related issues such as variations in head size and brain morphology (Lepage et al., this issue).

The pipeline features registration to the NMT v1.2 population template (Seidlitz et al., 2018), brain-masking, tissue classification, cortical surface reconstruction, and calculation of cortical morphometrics (thickness, surface area, and curvature). Surface-based registration to the NMT average surface allows for group analysis and regional analysis based on either the D99 or CHARM anatomical parcellations (Jung et al., this issue). Figure 5B illustrates a typical CIVET run with the preprocessing stages (top row: registration, brain-masking, tissue classification) and the generation of the cortical surfaces (bottom row). The preprocessing stages, which can be generated without cortical surfaces, can serve as the foundation for the analysis of other *in vivo* modalities (fMRI, qMRI, PET, EEG, MEG, sEEG, etc.) as well as *post-mortem* histological and genetic data. Images of the outputs from various viewpoints are provided for quality assessment.

The full pipeline was successfully run on the *in vivo* scans from the 31 subjects that were used to make the NMT and on the *ex vivo* scan of the D99 subject. The preprocessing stages of the pipeline were run on 95 PRIME-DE subjects from 15 sites that provided scans in the sphinx position with complete sex and age information (Milham et al., 2018). Efforts have also begun to further extend CIVET to the marmoset and, using archived data, the chimpanzee. The pipeline thus has the potential to make it possible to study surface-based morphometry across a range of primate species.

###### 3.2.2.5.2. NHP-Freesurfer

*Creating Freesurfer surfaces of macaque brain scans*

Freesurfer (Fischl, 2012) is one of the most widely used software packages for the creation of surfaces from volumetric MRI data in humans. While it can also be used to process non-human brains, its convenient recon-all pipeline that automates most of the processing steps cannot be readily applied to macaque brains due to its differences in size and contrast. NHP-Freesurfer offers a pipeline that does work for macaque brains. It is based on Freesurfer tools and a number of additional workarounds. All these steps are explained and pre-coded in annotated Jupyter Notebooks based on the bash-kernel that allow the user to run the code directly from the notebook. Unlike recon-all, which is mostly automated, creating high quality surfaces of macaque brains requires some manual adjustments that can be difficult to figure out for novice users. NHP-Freesurfer provides a full guide starting with one or more T1-weighted anatomical scans and ending with surfaces and flatmaps. To facilitate this process, NHP-Freesurfer uses freely available tools from the NMT template brain (Seidlitz et al., 2018), ANTs (Avants et al., 2008), and AFNI (Cox, 1996). In addition to the core Freesurfer functionality, NHP-Freesurfer also explains how to project volumetric statistical results to the Freesurfer surfaces with Freesurfer tools, but this workflow has been superseded by the NHP-pycortex package (see 3.2.3.5) that uses an adapted version of PyCortex (Gao et al., 2015) to obtain similar results with a much more flexible approach based on the Python programming language.

###### 3.2.2.5.3. Precon_all

*An automated Freesurfer and HCP compliant surface pipeline for preclinical animal models*

Precon_all is a surface reconstruction pipeline inspired by Freesurfer’s recon-all and adapted for use with non-human brains (Fischl, 2012). The great success of Freesrufer’s recon-all is in part due to the ease of use for the end user as well as the automation of surface generation from volumetric data. Precon_all aims to export this ease of use and automation to the animal imaging community. In order to achieve the flexibility that makes the precon_all pipeline work with a large range of species, a set of 5 easily drawn masks are required as input. Guidelines for drawing these masks are provided in the precon_all documentation. Once they are made, they can be used for an individual subject or added to the precon_all standards directory and used to automate surface reconstruction for a new species. Currently, precon_all comes preloaded with standards for the NMT v1.2 (Seidlitz et al., 2018) and a pig template.

Precon_all makes use of commonly used software packages, including FSL, ANTs, Freesurfer, and Connectome Workbench (Avants et al., 2008; Fischl, 2012; Glasser et al., 2013; Jenkinson et al., 2002). Written as a set of shell scripts, precon_all is called from the command line and, like Freesurfer’s recon-all, it can be run in stages for manual editing and correction of surfaces. These stages include brain extraction, segmentation via FAST or ANTs, cutting and filling, surface generation, and optional surface down-sampling. Outputs are contained within a subject directory similar to that of recon-all, allowing Freesurfer users to easily transition from human to animal imaging. Final outputs include Freesurfer and Human Connectome Project compatible cortical surface models, as well as all required NifTI images and transforms created in the surface reconstruction process. A set of scripts is provided to easily generate group average surfaces and registration templates which can then be used to create a custom FSaverage for individual species.

###### 3.2.2.5.4. PREEMACS

*Brain surfaces and cortical thickness from raw structural data*

PREEMACS (pipeline for PREprocessing and Extraction of the MACaque brain Surface) is a set of common tools, customized for Rhesus monkey brain surface extraction and cortical thickness analysis. Some of the advantages of using PREEMACS are: 1) it avoids manual correction, 2) the pipeline was developed in the standard MNI monkey space, 3) it provides visual reports in each module as quality control, and 4) its highly accurate surface extraction with vertex correspondence between subjects. PREEMACS has a modular design, with three modules running independently. These modules perform the canonical workflow for MRI preprocessing (Alfaro-Almagro et al., 2018; Esteban et al., 2019; Glasser et al., 2013) using different previously-available functions from FSL (Jenkinson et al., 2002), ANTs (Avants et al., 2008), MRtrix (Tournier et al., 2012), MRIqc (Esteban et al., 2017), DeepBrain (https://github.com/iitzco/deepbrain), and FreeSurfer (Fischl, 2012). Inputs to the pipeline should be one (or preferably more) T1- and T2- weighted volumes per animal from the same session. Module 1 prepares the raw volumes for initial preprocessing in six steps: volume orientation, conformation, image cropping, intensity non-uniformity correction, image averaging, resampling and skull-stripping. Module 2 is the quality control module. It was adapted from the MRI Quality Control tool (MRIqc) (Esteban et al., 2017) for humans to obtain image quality metrics and provide a visual report from the results of Module 1. Module 2 uses these quality control metrics to provide a classification of image quality that allows PREEMACS to estimate whether the input will yield an appropriate surface reconstruction. Finally, Module 3 obtains cortical thickness measures based on the brain surfaces using an NHP-customized version of Freesurfer v6. To evaluate the generalizability of this procedure, PREEMACS was tested on two different NHP datasets: PRIME-DE (Milham et al., 2018) (57 subjects) and INB-UNAM (5 subjects). Results (Figure 5C) showed accurate and robust automatic brain surface extraction for both datasets. PREEMACS thus offers a robust and efficient pipeline for the automatic NHP-MRI surface analysis.

#### 3.2.3. Functional MRI Tools

Functional MRI (fMRI) uses time-series data in an attempt to model the fluctuations in the imaging signal with some type of event structure (Figure 1C). These events can be external stimuli (i.e., task-based fMRI), or the time series of activity in another brain area (i.e., functional connectivity). Conceptually, the analysis of fMRI data is not radically different across species, but the checks required at each analysis step do differ. For example, animal scanner setups may produce different kinds of B0 inhomogeneity and distortions, due to customized coils or the use of other specialized equipment during acquisition. Both the potential use of anesthesia and contrast agents can furthermore impact processing choices such as hemodynamic response modeling and noise filtering. Importantly, pre-processing steps like motion correction and alignment to a high resolution anatomical scan or template brain often require different options and parameters compared to what one would choose for human data. For awake animals, body motion can cause distortions in the magnetic field resulting in slice-specific nonlinear deformations that can be especially difficult to correct. Proper head-fixation and extensive training will keep such problems to a minimum, but EPI distortion and brightness artifacts can still occur, requiring post-acquisition amelioration. The resources below contain several NHP specific solutions to deal with pre-processing and alignment correctly.

##### 3.2.3.1. afni_proc.py

*AFNI program that generates a complete fMRI processing script from a list of data files and the desired processing steps and options.*

AFNI’s afni_proc.py program (Cox, 1996) allows a researcher to create a complete fMRI processing pipeline for individual subjects, from raw inputs, through alignment to standard space, and regression modeling. To use it, one specifies input datasets (e.g., EPIs, anatomicals, tissue masks), major processing blocks (motion correction, warping to standard space, blurring, regression, etc.), and detailed choices for each block (e.g., what kind of alignment, the blur radius, order of the polynomial for baseline modeling and types of motion regressors). Thus, one can plan the processing hierarchically, for conceptual clarity and organization. The command generates a full, commented processing script, which is then executed to carry out single subject processing. In practice, a typical afni_proc.py command contains 20-25 options, which is a very compact way to specify a full pipeline.

For convenience and assured mathematical correctness, afni_proc.py automatically takes care of several aspects of the processing. For example, all warps (motion correction, B0 distortion, alignment to anatomical and to standard space) are concatenated before being applied to the EPI dataset to minimize smoothing due to regridding. This allows the user to focus on the parameter choices in processing, rather than on the technical programming aspects, which greatly reduces the number of bugs in an analysis stream, particularly if one updates or tweaks existing code. Because the processing script itself is explicitly created and saved, researchers can check exactly what steps are occurring in their analysis, and the whole process is documented, which increases reproducibility.

While afni_proc.py was mainly developed in conjunction with human brain researchers, it is fully compatible with animal data and has been applied in animal studies. For example, it integrates directly with AFNI’s @animal_warper command (see 3.2.2.4.2) and its list of possible hemodynamic response functions includes a stimulus response shape for MION (monocrystalline iron oxide nanoparticle), a contrast-agent that is commonly used in animal neuroimaging studies. When performing EPI-anatomical alignment, one can furthermore set the minimum “feature size” of structures to match to a value that is relevant for smaller animal brains.

There are currently two demos available in AFNI for macaque fMRI processing with afni_proc.py: one for task-based data (visual stimuli, MION contrast), and the other with resting state data from PRIME-DE (Milham et al., 2018). Each dataset uses the stereotaxic NMT v2 as its standard space and demonstrates nonlinear warp estimation with @animal_warper (see sections 3.2.1.1, 3.2.2.4.2, and Jung et al., this issue). AFNI is freely available, and most programs are written in C (as well as Python, R and shell), for generality and computational efficiency.

##### 3.2.3.2. C-PAC

*A flexible pipeline for performing preprocessing and connectivity analyses with various tools.*

The Configurable Pipeline for the Analysis of Connectomes (C-PAC) is an open-source platform that allows users to configure their own analysis pipeline for structural and functional MRI data. It is designed and tested for use with human, non-human primate, and rodent data. One of the key strengths of C-PAC is its ability to perform different processing strategies on the same dataset. Multiple tools can be specified for the same type of operation (for example, brain extraction or registration to a template), with the different results getting saved in separately-labeled output directories. This allows users to compare a set of methods and evaluate what type of preprocessing decisions may be best for their data.

A web-based pipeline editor is available (http://fcp-indi.github.io), but pipeline configuration files can also be edited with any text editor for quick and easy modifications. C-PAC also features a visual quality control interface which allows users to inspect the quality of brain extraction, segmentation, and registration outputs.

C-PAC is available as a Docker or Singularity container, allowing users to quickly get started. It is cloud-compatible through Amazon Web Services (AWS), and a machine image is available with a ready-to-use C-PAC Docker container for users who wish to run large-scale analyses. Finally, users can also point directly to a data directory that is hosted on the AWS cloud storage service (S3), and C-PAC will download and organize the data automatically.

##### 3.2.3.3. NeuroElf

*Versatile Matlab-based tool for visualization and (pre-)processing of fMRI data.*

Active development of NeuroElf has ceased since its developer is no longer active in neuroscience. This makes NeuroElf slightly outdated in some respects, as it relies on SPM8, an older version of the SPM package (Penny et al., 2004). Several components of the resource can however still be useful for the neuroimaging community. These components include the ability to import subject-level regression maps into a “GLM” format to rapidly test hypotheses and visualize bar and scatter plots of extracted regions, as well as some data export utilities, and its scripting capabilities.

##### 3.2.3.4. NHP-BIDS

*A BIDS-compatible Nipype-based pipeline for non-human primate fMRI data.*

NHP-BIDS is a pipeline for (pre-)processing of non-human primate fMRI data based on Nipype (Gorgolewski et al., 2011) and the BIDS (Gorgolewski et al., 2016) conventions for data storage. It is developed and maintained at the Netherlands Institute for Neuroscience (NIN). The accompanying Wiki-pages contain instructions to process data from scanner generated dicom images through to statistical results. The NHP-BIDS pipeline is available in fully configured form, set up for compatibility with the site where it was developed. However, NHP-BIDS is almost entirely written in Python and due to its implementation of the Nipype framework, it offers the user great flexibility in adapting the code-base to implement their own behavioral logging strategies or preferred choice of image processing modules. The first steps of NHP-BIDS deal with data-preparation and the creation of BIDS-compatible data structure. Shell scripts are offered that can easily be adapted to different data curation strategies to ensure automated data handling. Further data-processing steps are organized in Nipype workflows that can either be run on a local machine or offloaded to a cluster computing service. Instructions on how to use a SLURM-based job-scheduling system are included in the wiki.

The NHP-BIDS pipeline is modular and saves the intermediate results after every processing step for quality control. Standard modules used in the NIN-configuration are: 1) a minimal processing step that reorients the data to correct for the awake NHPs being scanned in sphinx position in a horizontal scanner; 2) a resampling step to ensure all data has isotropic voxels; 3) extensive preprocessing that includes non-rigid slice-by-slice realignment based on AFNI tools (Cox, 1996) and FSL-based motion correction (MCFLIRT) (Jenkinson et al., 2002); 4) registration to NMT template-space (Seidlitz et al., 2018); and 5) statistical analysis using FSL-FEAT (Woolrich et al., 2004, 2001). The resulting data can be visualized using standard software packages, or further processed for projection to the cortical surface using the packages NHP-Freesurfer and NHP-pycortex that are also made available by the NIN (see sections 3.2.2.5.2 and 3.2.3.5).

##### 3.2.3.5. NHP-pycortex

*Projecting volumetric statistical maps to macaque brain surfaces.*

Projecting fMRI activity dynamics or statistical maps on surface renderings of the cortex allows a much richer view of their spatial characteristics than can be obtained with static 2D slice image renderings. Creating surface renderings and projecting volumetric data to it can however be challenging, especially for non-human data. The Pycortex package (Gao et al., 2015) is a very flexible python-based toolbox designed to work with human brain surfaces that are generated with Freesurfer (Fischl, 2012). NHP-pycortex is an adapted version of the pycortex package that is compatible with the macaque brain surfaces generated with Freesurfer based tools (it is specifically tailored to the output of NHP-Freesurfer, see section 3.2.2.5.2). Jupyter Notebooks are provided to guide the user through the process of importing the Freesurfer surfaces into the Pycortex database and project data to it.

##### 3.2.3.6 Pypreclin

*A workflow pipeline dedicated to macaque functional and anatomical MRI preprocessing*

Pypreclin (Tasserie et al., 2020) was originally developed to face the challenges of artifacts induced by intracranial implants and body movements during awake fMRI acquisition. The pipeline development was further extended to a greater panel of experimental parameters such as BOLD-fMRI, CBV-fMRI (MION contrast agent), anesthesia with different pharmacological agents, or different RF coil configurations. Pypreclin is a Python module that is available on an open source repository (https://github.com/neurospin/pypreclin) with HTML documentation.

Pypreclin development aimed at including state-of-the-art algorithms to allow the automatic preprocessing of raw fMRI data. The diversity of acquisition conditions and hardware in the PRIME-DE database (Milham et al., 2018) was used to validate Pypreclin’s versatility and robustness across a large range of magnetic field strengths (1.5, 3, 4.7 and 7T), for data acquired with both single loop or multi-channel phased-array coils, with or without iron-oxide contrast agent, in awake and anesthetized animals, and both with or without intracranially implanted electrodes (Figure 6).

Compared to a previously used macaque fMRI preprocessing pipeline at the developer’s site (NeuroSpin Monkey), Pypreclin returned more accurate anatomical localization of neural activations in the gray matter in the awake state where body movements are often a major issue. The pipeline supports different brain templates for both *macaca mulatta* (Rhesus macaque) and *macaca fascicularis* (cynomolgus macaque) and can easily be customized, for instance with additional templates.

**Figure 6.**
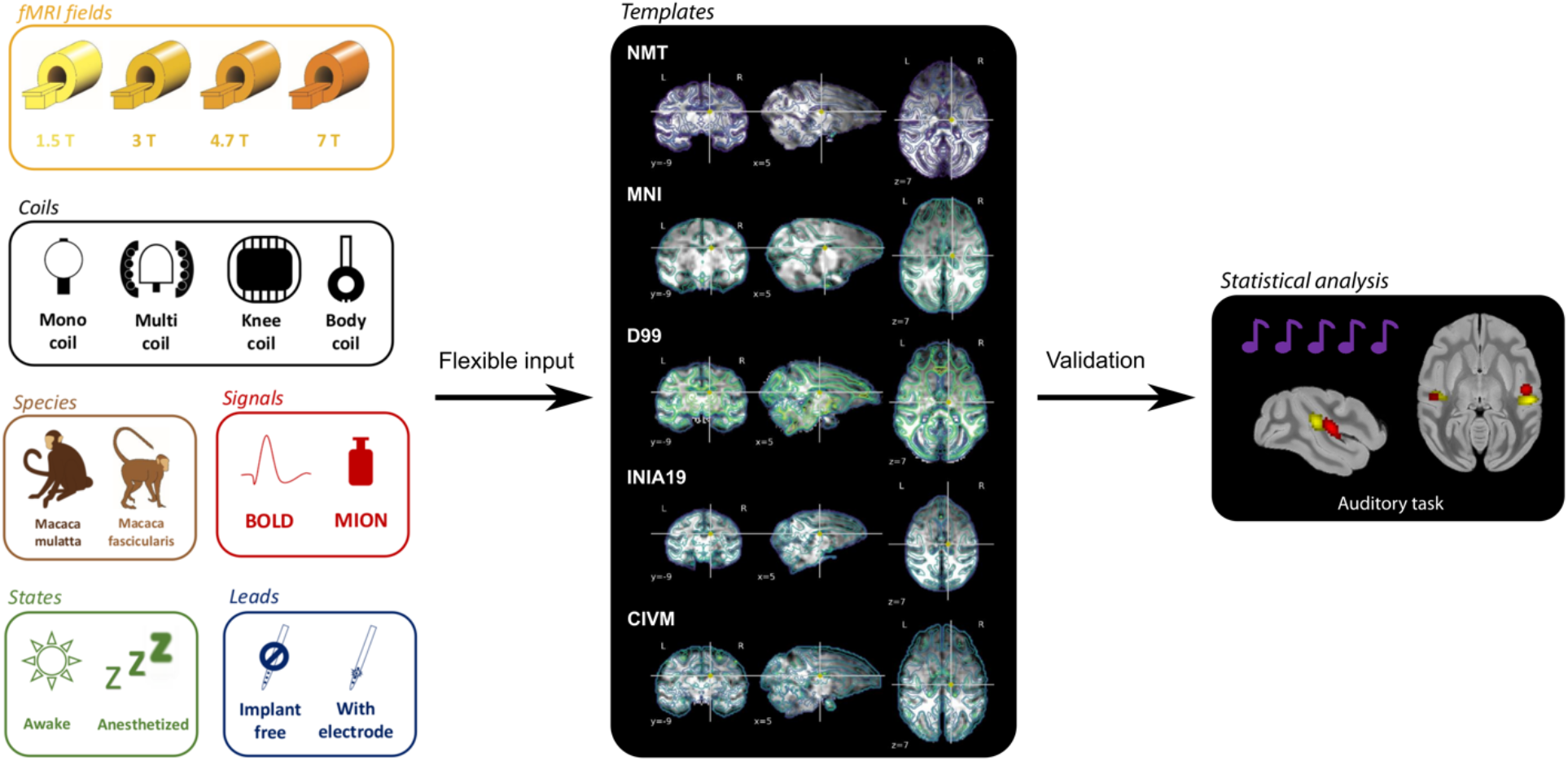
Functional MRI pipeline Pypreclin. The Pypreclin pipeline has been validated with a range of input data (left), a broad set of template brains (middle) and auditory task data. Preprocessing steps incorporated in Pypreclin include slice-timing correction, susceptibility distortions correction, motion correction, reorientation, bias field correction, brain extraction, normalization, coregistration and smoothing (for more details see: Tasserie et al., 2020).

#### 3.2.4. Diffusion-Weighted MRI Tools

Diffusion-weighted MRI is a technique to visualize the organization of the brain’s white matter structure (Figure 1D). Although tracer data are generally considered to be the gold standard for connectivity analysis, their invasive nature and high costs restrict their applicability. Diffusion MRI, however, can be performed both *ex vivo* and *in vivo* to obtain high quality, repeatable images that can be compared across species.

Several MRI software packages contain sets of diffusion processing tools that can be applied to either human or non-human datasets. The diffusion data processing and probabilistic tractography functions in FSL (Behrens et al., 2007; Jenkinson et al., 2002) were recently expanded with the Xtract tool to facilitate explicit comparison of fiber tracts across species. Xtract contains tractography libraries for humans and macaques derived from the same protocols (Warrington et al., 2020) and is currently being extended to include more species, including the chimpanzee (Bryant et al., 2020). AFNI (Cox, 1996) and TORTOISE (Pierpaoli, et al., 2010) have several diffusion processing tools and integrated features for distortion correction (Irfanoglu et al., 2015), deterministic and probabilistic tractography (Taylor and Saad, 2013), tensor-based morphometry (Hutchinson et al., 2018; Irfanoglu et al., 2016), and network-based structural analyses (Taylor et al., 2016b). Additionally, Dipy is a Python-based software toolbox for diffusion processing (Garyfallidis et al., 2014), and DSI-Studio contains tools for tracking and statistics with a particular focus on high angular resolution diffusion imaging (HARDI) techniques (http://dsi-studio.labsolver.org). These and other software solutions provide a wide range of functionalities that are often either complementary or integrable and facilitate analysis for various acquisition and study paradigms.

##### 3.2.4.1. Mr Cat

*Pipelines for processing of structural, functional, and diffusion non-human MRI data.*

The MR Comparative Anatomy Toolbox (Mr Cat; www.neuroecologylab.org) specializes predominantly in processing *post-mortem* data of a wide variety of brains, which are preprocessed using adaptations of FSL tools implemented in the ‘phoenix’ module. A number of post-processing modules are furthermore aimed at providing quantitative comparisons of brain organization across species. These are often based on connectivity measures, including matching of areal connectivity fingerprints across species (Mars et al., 2016) or comparisons of ‘connectivity blueprints’ of the cortex with the whole brain that allow the description of different brains in terms of a common connectivity space (Mars et al., 2018b). Recent extensions of this approach focus on the comparison of brain organization measured by multiple modalities, testing in effect whether the relationship between distinct aspects of brain organization differs across species (Eichert et al., 2020).

##### 3.2.4.2. Diffusion-MRI

*Pre- and postprocessing steps for diffusion-weighted imaging*

The Diffusion-MRI repository hosts Jupyter notebooks in Python and bash that guide users through multiple pre- and post-processing steps in the analysis of diffusion-weighted imaging (DWI) data. A step-by-step example analysis is provided of macaque DWI data from PRIME-DE (Milham et al., 2018). While the workflows were developed and tested with macaque data, they can easily be adapted for other primate brains. The workflow, which is based on tools from the Nipype (Gorgolewski et al., 2011), Dipy (Garyfallidis et al., 2014), FSL (Jenkinson et al., 2012) and MRtrix3 (Tournier et al., 2012) software libraries, requires single-shell or multi-shell DWI data in NIfTI file format as input, with reverse phase-encoding acquisition containing at least one non-diffusion-weighted image. Preprocessing includes denoising (Veraart et al., 2016), correction for susceptibility distortions, eddy current distortions, and subject movement artifacts using TOPUP (Andersson et al., 2003) and EDDY (Andersson and Sotiropoulos, 2016). Post-processing includes standard and advanced DWI models to fit the diffusion signal, including Diffusion Tensor Imaging (DTI) (Basser et al., 1994), Diffusion Kurtosis Imaging (DKI) (Jensen et al., 2005), Neurite Orientation Dispersion and Density Imaging (NODDI) (Zhang et al., 2012), Single-Shell 3-tissue Constrained Spherical Deconvolution (SS3T-CSD) (https://3tissue.github.io), and Multi-Shell Multi-Tissue Constrained Spherical Deconvolution (MSMT-CSD) (Jeurissen et al., 2014). The outputs are parametric maps extracted from the various model fits. The DTI model results in maps of axial, radial, and mean diffusivity, as well as fractional anisotropy. The DKI model – an extension of the DTI that captures diffusion non-gaussianity – also outputs axial, radial, and mean kurtosis. The three-compartment NODDI is fitted using Accelerated Microstructure Imaging via Convex Optimization (AMICO) (Daducci et al., 2015) and computes maps of Intra-cellular Volume Fraction (a measure of neurite density), cerebrospinal fluid (CSF) volume fraction, and fiber orientation dispersion. SS3T- and MSMT-CSD are used to estimate the multi-tissue orientation distribution function followed by whole-brain tractography (Figure 7A).

**Figure 7.**
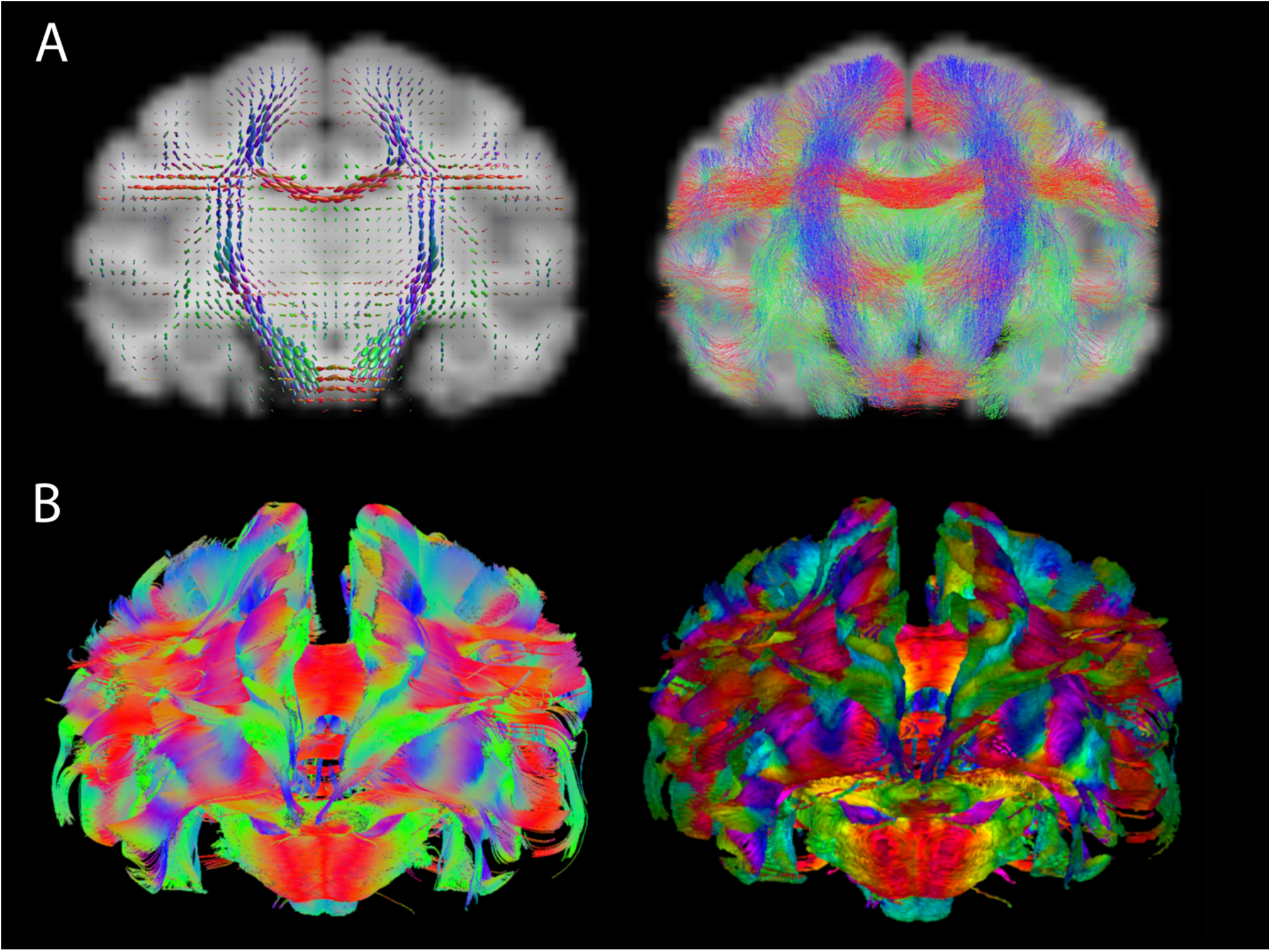
Diffusion-weighted MRI. **A**) Result from the Diffusion-MRI resource using macaque data. Fiber orientation distributions (left) and tractography (right). **B**) Examples of DTI-based tractographic output in a high-resolution macaque dataset in AFNI, following distortion correction with TORTOISE. Both images show a (frontal) coronal view of whole brain tracking using AFNI’s 3dTrackID: the left panel shows the results of deterministic tracking (as tract fibers), and the right panel shows the results of probabilistic tracking (as WM volume surfaces). Coloration reflects the orientation of the local structure relative to the coordinate frame, where red, blue and green are parallel to the x-, y- and z-axes, respectively. In the probabilistic panel, the FA value modulates the brightness (higher FA is brighter). The images are displayed using SUMA.

##### 3.2.4.3. FATCAT and TORTOISE

*DWI processing, distortion correction, tractography and various group analyses*

Diffusion-based imaging provides a great deal of structural information about the brain, with a particular emphasis on WM properties. It also presents particular processing and quality control (QC) challenges for limiting the effects of distortions due to subject motion, eddy currents and EPI inhomogeneity. AFNI’s FATCAT (Taylor and Saad, 2013) and the TORTOISE toolbox (Pierpaoli, et al., 2010) provide a full set of programs for processing and visualizing diffusion data, from DICOM conversion through distortion correction to group analysis, including quality control. FATCAT contains validated deterministic and probabilistic tracking tools (Taylor et al., 2012) that can be used for DTI- and HARDI-based modeling. Examples of tractographic output for a high-resolution macaque dataset are shown in Figure 7B. These tools can be useful in performing network-based comparisons of structural properties (Taylor et al., 2016b), possibly in conjunction with functional network studies. TORTOISE provides methods for group analysis based on separate comparisons of structural properties from diffeomorphic registrations (DR_TAMAS; (Irfanoglu et al., 2016). Study design is important for all aspects of analysis, but the DWI acquisition method particularly affects how well distortion artifacts can be reduced. Acquiring sets of DWIs with opposite phase encoding has been shown important in reducing EPI inhomogeneity distortions (Irfanoglu et al., 2019, 2015), especially in the presence of subject motion (Taylor et al., 2016a). A T2-weighted structural volume with fat suppression can furthermore act as a useful reference for further reduction of geometric distortions. TORTOISE’s tools have been designed to take advantage of such acquisitions.

Multiple demos for TORTOISE and AFNI’s FATCAT are available. The FATCAT_DEMO2 illustrates full single subject processing of a dual-phase encoded DWI dataset, including probabilistic tractography on gray matter parcellation. Each step of processing automatically generates QC images to help evaluate the data. These include alignment images, subject motion plots, directionally encoded color (DEC) maps, thresholded FA maps, and ROI overlays. The FAT_MVM_DEMO provides an example of combining FATCAT tractography results from a network of target ROIs with multivariate modeling (MVM) for a hierarchical group analysis: first, at the “network level” with an omnibus F-statistic, and then zooming in at the “ROI level” with post hoc t-tests. The techniques in each of these demos can be applied directly to other human or animal studies.

#### 3.2.5. Data sharing

As an extension of the PRIME-DE consortium, which was established to openly share NHP neuroimaging data (Milham et al., 2020, 2018), PRIME-RE also maintains a list of NHP neuroimaging data-sharing initiatives. Similar to the analytical resources described above, anyone can submit a (link to a) data resource for inclusion on PRIME-RE using a simple submission form. The data sharing section of PRIME-RE currently lists PRIME-DE, OpenNeuro, and NeuroVault. The PRIME-DE initiative (Milham et al., 2018) specifically focuses on NHP data and comprises data from 22 different sites all over the world. The database contains structural, functional and diffusion data; BOLD and contrast-agent data; awake and anesthetized scans; resting-state and task-based fMRI. OpenNeuro, previously known as OpenfMRI (Poldrack et al., 2013; Poldrack and Gorgolewski, 2017) hosts MRI, MEG, EEG, iEEG, and ECoG data. Most of the data are from human subjects, but OpenNeuro is not restricted to human data and some NHP data-sets are now available as well. Neurovault (Gorgolewski et al., 2015) specializes in sharing un-thresholded statistical maps, mask files, parcellation maps, and any other voxelwise data. Originally restricted to human imaging data, the framework was recently extended to allow non-human data as well (Fox et al., this issue). For NHP data, volumetric data should be registered to the NMT, a processing step that can easily be done using several of the tools listed on PRIME-RE (see section 3.2.2).

## 4. Discussion

Neuroimaging with non-human primates requires highly specific experimental and analytical expertise. Recent initiatives such as PRIME-DE (Milham et al., 2018) have made a wealth of shared NHP neuroimaging data openly available, but the analysis of these datasets demands appropriate analytical expertise, tools, and workflows. To facilitate the study of NHP neuroimaging data, PRIME-RE was established as an infrastructure for knowledge and resource sharing, collaboration and communication. The PRIME-RE website serves as a central knowledge hub for the NHP neuroimaging community. It is home to a structured dynamic overview of relevant analytical resources, and a continuously evolving wiki that describes the challenges and potential solutions of all facets of NHP neuroimaging. Both these knowledge bases are collectively curated by community-driven practices. This content makes PRIME-RE a NHP-specific complement to more broadly oriented sites, such as the Neuroimaging Tools and Resources Collaboratory (NITRC, https://www.nitrc.org) (Kennedy et al., 2016; Luo et al., 2009) that catalogues data, tools and pipelines for all general aspects of neuroimaging. A common data structure to reference research resources, such as the RRID (Research Resource Identifier) (Bandrowski et al., 2016) might allow seamless integration of the resources in PRIME-RE and NITRC in the future.

It is common for development teams of broadly used neuroimaging packages to organize courses focused on the use of their specific package (e.g., AFNI^1^, FSL^2^, or SPM^3^). In recent years, open science initiatives such as Brainhack (Craddock et al., 2016) and the NeuroHackademy, have led to a boost in the creation of tutorials, courses, demos and workshops that more generally cover different aspects of neuroimaging and open science practices. While this content is often spread over various websites, some efforts are being made to curate this information (e.g, https://learn-neuroimaging.github.io/tutorials-and-resources). Besides its community-curated wiki and links to tutorials and documentation for individual resources, PRIME-RE also hosts a list of links to external tutorial and resource collections that may be useful for the NHP neuroimaging community.

The PRIME-DE/RE initiatives follow on prior large-scale human neuroimaging resources, such as the 1000 Functional Connectomes Project (Biswal et al., 2010), ADHD-200 (Brown et al., 2012) or ABIDE (Di Martino et al., 2014). In these projects, data sharing was complemented by open source pipelines for preprocessing and analyzing the data (Configurable Pipeline for the Analysis of Connectomes) (Craddock et al., 2013). While these accompanying tools may be limited in resolving all potential issues, their early release lowers the barrier for users to engage with the shared data, and accelerates the development of novel analysis solutions. By focusing on community building, PRIME-RE aims to provide a broad dynamic framework to channel the efforts of the NHP neuroimaging community and evolve together. In the future, PRIME-RE may be expanded to include an even broader spectrum of resources involved in NHP neuroimaging. One can think of blueprints for hardware (e.g., chairs, behavioral response devices, and coils), electronic schematics, models for 3-D printing or laser cutting; protocols for data acquisition (e.g., sequences), animal care, housing, training and handling. Additional resource categories can be created for protocols and tools targeted at the combination of NHP neuroimaging with other experimental approaches such as brain perturbation techniques (Klink et al., this issue), electrophysiological recordings, and histology. The Wiki could furthermore be expanded to address task or experimental design, validation approaches, quality control, and different types of univariate and multivariate analysis methods, although some of these are not specific to NHP neuroimaging.

PRIME-RE supports best practices of open science at every step from data acquisition, through data organization, to code structure and analysis. Contributors and users are encouraged to adhere to the FAIR principles (Wilkinson et al., 2016) and both communicate and document their tools and data. Compared to human neuroimaging, NHP neuroimaging has been lagging behind in adopting such collaborative open science initiatives, and we can only speculate about the reasons why. It is conceivable, for instance, that due to the large amount of work involved in overcoming the challenges of NHP neuroimaging, the researchers involved might be a bit more protective of their solutions. In many cases, these solutions are furthermore developed by early career researchers that tend to get judged on their research output and not necessarily on the tools they develop for use by others in the field. Systems neuroscience and translational neuroimaging are strongly multidisciplinary in nature, combining elements of biology, psychology, physics, engineering and more. Researchers involved in the field cannot be expected to be an expert in all these disciplines and a substantial proportion of them may not have extensive formal training in mathematics, physics, or programming which could make them reluctant to share their custom-written code for outside scrutiny. Another possibility is that such sharing initiatives do actually exist but that they are rather difficult to find for outsiders. For instance, because they reside on a laboratory’s internal servers or personal websites. However, people’s attitudes towards openly sharing the resource solutions they have developed appear to be improving (Balbastre et al., 2017; Tasserie et al., 2020) and new developments in information technology promote such initiatives while simultaneously creating avenues to assign explicit credit to developers.

A collaborative open approach to NHP neuroimaging will not only foster collaboration in the short term, but also guide future development efforts, enhance reproducibility, and benefit the whole community in the long term. The contribution template for PRIME-RE requires a minimum set of metadata to ensure a consistent and comprehensive resource index (see Table 1). The website provides a graphic interface that simplifies database navigation and makes it easy for researchers to discover relevant tools, data, and learning material. PRIME-RE’s agile governance structure is decentralized and open (hosted on GitHub) with clear guides to help researchers integrate into the community and contribute to it. The original resources listed on PRIME-RE are maintained by their respective developers and indexed in a completely community-driven way. Acting as a community-based curation layer on top of open resources reduces processing bottlenecks that could be encountered if a small single team were responsible for maintaining and updating the system, while simultaneously facilitating scalability, and inclusiveness.

The maintenance of experimental and analytical code is a challenge in academic research. The adoption of code development best practices, such as version control, comment inclusion, testing, coding style, and continuous integration (Eglen et al., 2017), accelerates the continuous improvement of tools and assists in the swift detection and fixing of bugs. Unfortunately, these practices are not yet common in academia. We strongly advocate for their adoption and are hopeful that open and informal communication between researchers and developers at all levels of NHP neuroimaging research, as facilitated by PRIME-RE, can accelerate this process.

What makes a neuroimaging resource suitable for use with NHPs or other non-human species? While toolboxes that were primarily developed for use in humans sometimes generalize to, or contain options for use, in non-human imaging, more often such packages require substantial customization for use in non-human species. Built-in assumptions that, while valid for most human neuroimaging data, are not met by animal data include subject orientation in the scanner, signal quality (contrast), brain size, and field-of-view. An open conversation about such limitations between developers and researchers could help the development of tools that are innately capable of processing data from a range of different species (and developmental stages). Such tools are invaluable for cross-species comparisons that inform our understanding of brain evolution and development. Software customizations that are often required to make a package compatible with NHP data can make it difficult to assess data quality and intermediate results of analytical steps in more comprehensive pipelines. Such quality control is underutilized in neuroimaging in general but it is even more crucial for non-human data that is analyzed with tools that were not developed for this type of data or extensively tested on it.

Finally, because NHP-specific analysis tools are often developed by a single team for one particular research environment, local dependencies could make it difficult for research teams elsewhere to employ such tools in their own research, even if they are openly shared. The increasing popularity of modular pipelines (Gorgolewski et al., 2011; Mourik et al., 2018; Tasserie et al., 2020) is an important improvement for cross-site compatibility and reproducibility and some of the broader software packages on PRIME-RE contain functionality that specifically addresses this challenge. Packages like AFNI have modularity built-in, while others, like FSL, integrate with ‘notebook-style’ scripting modules. A wealth of Python-based tools developed by the NIPY community (https://nipy.org) furthermore provides access to functions from many different analytical toolboxes through standardized wrapper modules that can be used as building blocks for analysis and visualization pipelines. The sharing of tools in pre-packaged containers that include all dependencies (e.g., Docker or Singularity images) can also make it easier for users to try a particular piece of software. Standardization of tools and quality assessment methods benefit from predictable data structures and file formats. While PRIME-RE does not host any data itself, it does promote the use of standardized data structures like BIDS and platform-neutral data formats (e.g. NifTI/GifTI/CifTI).

## 5. Conclusion

With PRIME-RE, we introduce a collaborative platform to address the experimental and analysis needs of the NHP neuroimaging research community. By allowing the community to curate relevant resources, disseminating and encouraging open science practices, and strengthening communication and interactions among researchers and developers, we aim to accelerate reproducible discovery, minimize redundant efforts, and maximize efficiency of this invaluable form of translational and comparative neuroscience.

## CRediT authorship contribution statement

AM, PCK: Conceptualization, Methodology, Project administration, Data curation, Writing, Software, Resources, Visualization, Supervision.

NS, JuS, BC, KH: Conceptualization, Writing, Software, Visualization

DSM: Writing, Project administration, Supervision

KKL, RM, TX, DG, BJ, JaS, PT, RT, EAG-V, CS, XW, RAB, RD, HCE, PG-S, SG, RH, ClL, CiL,

PM, HM, MM, MGPR, JT, LU: Writing, Software, Visualization

## Code and data availability

Both the code for the PRIME-RE website (https://prime-re.github.io) and its Wiki pages are available on GitHub (https://github.com/PRIME-RE/prime-re.github.io). All resources described in this paper and listed on PRIME-RE are freely available from their respective developers or maintainers. Every listing on PRIME-RE comes with a link to the actual resource and a brief statement on possible usage restrictions (e.g., citing a specific paper). The list of more common software packages may contain packages that require users to purchase a license, but most are freely available as well.

## Acknowledgements

We would like to thank Patrick Markwalter for assistance with compiling references, editing and formatting. We thank the following people for their contributions to shared resources: Richard C. Reynolds, Gang Chen, and Robert Cox (AFNI, esp. afni_proc.py); Konrad Wagstyl and Alan C. Evans (CIVET); Leslie Ungerleider (CIVET, NMT); Cameron Craddock (C-PAC); David Meunier and Régis Trapeau (Macapype); Jochen Weber (NeuroElf); Jonathan Williford (NHP-BIDS); Borja Ibañez (precon_all); Arun Garimella, Felipe Mendez, and Luis Concha (PREEMACS); Antoine Grigis and Béchir Jarraya (Pypreclin); Gabriel Devenyi, Nikos K. Logothetis, and George Paxinos (SARM); Afonso Silva and David Leopold (MBM); Lennart Verhagen and members of the Cognitive Neuroecology Lab (MrCat). This research was supported in part by the Intramural Research Program of the NIMH and utilized the computational resources of the NIH HPC Biowulf cluster (http://hpc.nih.gov). Pypreclin work was supported by the Fondation pour la Recherche Medicale (FRM grant number ECO20160736100 to JT), Fondation de France, Human Brain Project (Corticity project). RT and KH are supported by ANR-19-DATA-0025-01 NeuroWebLab. RBM is supported by the Biotechnology and Biological Sciences Research Council (BBSRC) UK [BB/N019814/1]. The Wellcome Centre for Integrative Neuroimaging is supported by core funding from the Wellcome Trust [203129/Z/16/Z].

## Footnotes

1 https://afni.nimh.nih.gov/pub/dist/doc/htmldoc/educational/bootcamp_recordings.html

2 https://fsl.fmrib.ox.ac.uk/fslcourse

3 https://www.fil.ion.ucl.ac.uk/spm/course

